# Conserved and species-specific transcriptional responses to daily programmed resistance exercise in rat and mouse

**DOI:** 10.1101/2023.08.23.552047

**Authors:** Mark R Viggars, Hazel Sutherland, Christopher P Cardozo, Jonathan C Jarvis

**Affiliations:** Research Institute for Sport & Exercise Science, Liverpool John Moores University, Liverpool, United Kingdom; Department of Physiology and Aging, University of Florida, Gainesville, Florida United States; Myology Institute, University of Florida, Gainesville, Florida, United States; James J Peters VA Medical Center, Bronx, New York, and Icahn School of Medicine at Mount Sinai, New York, NY, United States

**Keywords:** Muscle hypertrophy, transcription, exercise, metabolism, functional electrical stimulation, muscle atrophy

## Abstract

Mice are often used in gain or loss of function studies to understand how genes regulate metabolism and adaptation to exercise in skeletal muscle. Once-daily resistance training with electrical nerve stimulation produces hypertrophy of the dorsiflexors in rat, but not in mouse. Using implantable pulse generators, we assessed the acute transcriptional response (1-hour post exercise) after 2, 10 and 20 days of training in free living mice and rats using identical nerve stimulation paradigms. RNA-sequencing revealed strong concordance in the timecourse of many transcriptional responses in the tibialis anterior muscles of both species including responses related to ‘stress responses/immediate-early genes’, and ‘collagen homeostasis’, ‘ribosomal subunits’, ‘autophagy’ and ‘focal adhesion’. However, pathways associated with energy metabolism including ‘carbon metabolism’, ‘oxidative phosphorylation’, ‘mitochondrial translation’, ‘propanoate metabolism’ and ‘valine, leucine and isoleucine degradation’ were oppositely regulated between species. These pathways were suppressed in the rat but upregulated in the mouse. Our transcriptional analysis suggests that although many pathways associated with growth show remarkable similarities between species, absence of an actual growth response in the mouse may be because the mouse prioritises energy metabolism, specifically replenishment of fuel stores and intermediate metabolites.

## Introduction

Muscle strength has been identified as a predictor of all-cause mortality, so interventions that improve or ameliorate the deterioration of muscle strength with age or disease are important preventative and interventional aspects of medicine^1,2^. Indeed, resistance exercise training has been shown to be beneficial in promoting health in a variety of disease states including ageing, metabolic disorders, cardiovascular disease, cancer cachexia and neurological conditions^3–6^.

Resistance exercise causes metabolic and mechanical perturbations in the muscle that initiate acute signaling cascades^7–11^, with both local and systemic effects^12^. With regular resistance training, these transcriptional changes can lead to increases in muscle size, quality, strength and improvements in metabolic function, but the molecular underpinnings of such adaptations are only partially understood.

Mice remain in many cases the most desirable model for research into the molecular biology underpinning the benefits of exercise because the mouse genome is well annotated, and because tools exist for relatively easy and precise manipulation of the genome so that gene function can be investigated by gain or loss of function studies with global or tissue specific techniques^13^. Adaptation to endurance exercise can be easily studied in rodents through voluntary or forced wheel/treadmill running, but interventions that mimic resistance-like exercise are more challenging to set up^14^. An established method to mimic the ‘sets and reps’ style of resistance-like exercise is the use of high frequency contractions delivered to the hindlimb via electrical stimulation of the sciatic nerve so that the ankle dorsiflexors is resisted by simultaneous contractions of the plantarflexors. In the rat, this results in extensive hypertrophy of the dorsiflexors (10-30%) in a matter of weeks^15–19^, without extensive damage or regeneration. In recent years, we have developed this co-contraction technique by means of implantable stimulators for use in free living rats and mice so that anesthesia during stimulation is not required. Similar exercise paradigms in the mouse seem to be protective against muscle wasting^20,21^, but to the best of our knowledge result in little or no muscle growth^22^ and we confirm that finding here over a period of 3 weeks of daily training. Several papers on the other hand have used the co-contraction paradigm to interrogate the signalling response to high force contractions in mice^11,23–25^.

There are a number of potential molecular factors that may determine these differences in muscle growth between species, including but not limited to the basal transcriptional, epigenetic and metabolic properties of the hindlimb muscles that dictate the responsiveness to exercise and protein turnover rates.

Here we focus on the transcriptional differences that are associated with the lack of increase in muscle mass in mouse dorsiflexors. We used RNA-sequencing to compare the acute transcriptional responses to resistance exercise as the training status changes over a period of three weeks of daily training in mice and rats. Uniquely, our resistance exercise paradigm allows identical nerve stimulation patterns to be delivered by implantable pulse generators in free living mice and rats providing a highly controlled contraction paradigm. Our transcriptome analysis highlights both concordant or conserved and discordant or species-specific transcriptional responses to the identical nerve-stimulated resistance exercise over a period of daily training which may underpin the differences in growth response.

## Materials and Methods

### Experimental Design

The animal experiments were conducted under the provisions of the Animals (Scientific Procedures) Act 1986 and approved by the British Home Office (PA6930221).

### Resistance Training Protocol & Pattern

Animals received 1 session per day of SpillOver resistance training in the left hind-limb via stimulation from an implanted pulse generator (IPG) as previously described^17–19,26^, for 2, 10 or 20 days. Training was conducted between ZT1-4, where ZT0 indicates lights on and ZT12 indicates lights off. The dorsiflexor muscles received supramaximal activation via a cathode placed underneath the common peroneal nerve, while the anode was positioned underneath the tibial nerve and stimulated the plantarflexor muscles with the stimulus current adjusted during the implant operation to provide enough recruitment to provide resistance against the contraction of the dorsiflexors so that the dorsiflexors contracted isometrically or with slightly eccentric contractions for the duration of the study.

Daily training was delivered automatically by the IPG and consisted of an initial 10 seconds of preparatory stimulation at a low frequency (F = 4 Hz, phase width = 258 µs, current = approximately 1 mA), followed by 5 sets of 10 tetanic contractions at 100 Hz. Each contraction lasted for 2 s with 2 s rest between contractions and 2.5 minutes of rest between sets. Stimulation was delivered only in the left hind-limb. Muscles of the right hind-limb acted as unstimulated contralateral controls.

### Implantable Pulse Generators

Silicone encapsulated radio frequency controlled implantable pulse generators (IPGs) (MiniVStim 12B, Competence Team for Implanted Devices, Center for Medical Physics and Biomedical Engineering, Medical University Vienna, Austria) were used to deliver electrical impulses which invoke contraction of the targeted nerves/muscles.

Pulse generators were programmed wirelessly via an external programming device and an Android driven tablet computer (Xperia Tablet Z, Sony Corporation, Tokyo, Japan) as previously described^17^. This allowed for fine adjustment of the stimulation amplitude individually for each animal. After initial setup, the customized stimulation pattern was saved to the implant, enabling automatic daily delivery of SpillOver stimulation to the targeted nerves. The electronic circuit was connected for use in rats to a 1/3N lithium battery. In mice, the same circuit was driven by a 1220 lithium battery to reduce the overall size of the implant.

### Surgical Procedure

Animals were anaesthetised during implant procedures by inhalation of a gaseous mixture of isoflurane in oxygen at approximately 3% for induction and 1-2% for maintenance. Once anaesthetised, a subcutaneous injection of Enrofloxacin for antibiotic cover (5mg/kg-1 body mass (Baytril®) and an intramuscular injection of Buprenorphine for analgesia at 0.05mg/kg body mass in rat and 0.1 mg/kg body mass in mouse (Temgesic, Indivior, Slough, UK) into the right quadriceps were given. Strict asepsis was maintained throughout the procedure. In rats, the IPGs were implanted into the abdominal cavity accessed by a lateral incision through the skin and peritoneum, between the rib cage and pelvis on the left side of the animal. A polyester mesh attached to the IPG was incorporated into the suture line closing the peritoneum, securing the device against the abdominal wall. In the mouse, the implant was placed subcutaneously on the flank. Two PVC-insulated stainless-steel electrode leads (Cooner Sales Company, Chatsworth, California, USA) with terminal conductive loops, were fed from the implant site and tunnelled under the skin to the lateral side of the left hind-limb, just above the knee. A second incision was made through the skin and biceps femoris muscle to give access to the common peroneal nerve (CPN) under which the cathode was placed (to stimulate the dorsiflexors). The anode was placed in the muscular tissue deep to the tibial nerve about 5 mm distal to its bifurcation from the sciatic nerve to allow SpillOver stimulation to produce additional partial activation of the plantarflexors and thus to resist the contraction of the dorsiflexors. All incisions were closed in layers and 7 days were allowed for recovery from surgery before the start of the training protocol which lasted for 2, 10 or 20 days.

### Histological Analysis

Fiber cross-sectional area labelling for dystrophin, NADH activity and haematoxylin and eosin staining, and subsequent analysis was performed as previously described^17,19,26–28^. Mouse tibialis anterior samples were sectioned at 10μm using an OTF5000 Cryostat (Bright Instruments, UK) onto Thermo Scientific SuperFrost Plus Adhesion slides (Thermo Fisher Scientific Inc., Waltham, MA). Muscle cross sections were labeled with a polyclonal dystrophin antibody (Cat. No. PA5-32388; Thermo Fisher Scientific) at (1:200) and appropriate fluorescently conjugated anti-rabbit IgG (H + L) secondary with Alexa Fluor 488 (1:500) was used to demarcate the inside of the sarcolemma.

Once labelled, whole muscle cross sections were imaged using a Zeiss LSM 900 confocal with a x20 objective for fluorescence and a Leica LMD6 microscope with a x10 objective for brightfield imaging. Multiple images were automatically stitched together using the tilescan feature in the Zen Blue v 3.8 system for fluorescence and the Leica Application Suite for brightfield. Images were automatically processed for fiber cross-sectional analysis using MyoVision 2.0 software using default parameters for mouse^26^.

### Muscle Sampling, RNA Isolation and RNA Library Preparation & Sequencing

Animals were euthanised using a rising concentration of carbon dioxide, followed by cervical dislocation. Tibialis anterior (TA) muscles from both hind limbs were immediately harvested, cleaned of excess connective tissue, and weighed. A small cross-sectional sample was frozen in isopentane above liquid nitrogen for histology and the remainder flash frozen in liquid nitrogen for subsequent RNA-extraction. About 30mg of mouse TA muscle was added to MagNA Lyser Green Bead tubes (Roche 03358941001) with 600ul lysis buffer (PureLink RNA Mini Kit, Invitrogen 12183018A) and homogenised using a MagNA Lyser (Roche) at 6500rpm for three, 40 second sets with cooling in a rotor block at −20°c in between. Homogenates were centrifuged at 12000g for 10 minutes. Supernatant (400ul) was mixed with 400ul ethanol and added to spin cartridge (PureLink RNA Mini Kit). RNA was bound to the column by centrifugation and was washed multiple times per the manufacturer’s recommendations. RNA was eluted in nuclease free water from the column. RNA integrity numbers (RIN) were determined on an Agilent TapeStation 2200 (Agilent, CA, USA). All samples prepared for sequencing had RIN values of at least 8. Libraries were created using Poly-A tail selection using the Illumina mRNA Prep kit (Illumina, Inc. CA, USA) kit and pooled to equal molarities and sequenced on the Novaseq 6000 (Illumina, Inc. CA, USA) S4 2×150bp platform by the University of Florida Interdisciplinary Centre for Biotechnology Research (ICBR), RRID:SCR_019145 and SCR_019152. RNA-sequencing from the rat experiments was taken from the GEO accession GSE196147 but realigned to the latest genome annotation model as described below. The RNA extraction and sequencing from the rat tissue has been described previously^17^.

### Bioinformatic Analysis

FastQ files were imported to Partek® Flow® Genomic Analysis Software Partek Inc. (Missouri, USA) for pipeline processing. Pre-alignment QA/QC was performed on all reads prior to read trimming below a Phred quality score of 25 and adapter trimming. STAR alignment 4.4.1d was used to align reads to the *Rattus Norvegicus*, Rn7 genome assembly or the *Mus Musculus*, mm10 genome assembly^29^. The data generated for the rat has previously been published with analysis and alignment performed on the earlier Rn6 genome assembly. Rn7 has significant improvements from Rn6 including increased genome coverage and reduced gaps between scaffolds making it comparable in quality to the current human and mice genome assemblies^30,31^. Aligned reads were quantified to Ensembl annotation models and gene expression normalised using DESeq2 median of ratios^32^ and DEGs identified through the Partek® Flow® DESeq2’s binomial generalized linear model with Wald testing for significance between conditions set at an FDR of 5%. QA/QC data and normalized count information is available in Supplementary File 1.

Filtered lists for common and species specific differentially expressed genes (DEGs) were extracted and imported to DAVID. DAVID functional annotation was performed using a high stringency cut-off for pathways associated with the KEGG pathway database, the Reactome pathway database and the Gene Ontology (GO) biological process and cellular component databases^33^. Data contained in individual gene plots was generated by Log^2^ transforming the Z-scores of normalised Deseq2 counts.

### Statistical information

Muscle mass data are presented as the % change between the left experimental hind-limb and right internal contralateral control hind-limb for normalised muscle mass (mg/kg bodyweight). The resultant percentage changes were then compared via one-way ANOVA, followed by Tukey’s post-hoc analysis to confirm differences between groups in GraphPad Prism 9.0 software. All data are presented as mean ± standard deviation (SD). Pearson’s correlations were also performed in GraphPad Prism 9.0 software. Individual gene plots are presented as Log^2^ Z-scores of normalised Deseq2 counts.

## Results

### The timecourse of changes in muscle mass following identical activity/loading patterns in mice and rats

Daily SpillOver resistance training in rats results in a progressive increase in muscle mass after 2 (4.2 ± 3.3 %, *P*= 0.37), 10 (13.1 ± 5.2 %, *P*< 0.0001), and 20 days of consecutive resistance exercise (16.1 ± 4.2 %, *P*< 0.0001) compared with sham surgery (−0.9 ± 1.4 %), as previously reported in a separate study^19^. The increase in muscle mass plateaus between 10 and 20 days of training (*P* = 0.23), (Figure 1B). Similar experimental paradigms in other laboratories have yielded similar results in the rat^15,34^. Interestingly, the same nerve stimulation paradigm in mice has anecdotally been reported not to result in a hypertrophic response. In our mouse experiments, we observed small reductions in muscle mass after 2 (−6.4 ± 0.5 %), 10 (−3.8 ± 5.6 %), and 20 days (−8.1 ± 4.4 %) of training respectively despite using the identical nerve stimulation paradigm as used in the rats (Figure 1B). Through immunofluorescent dystrophin labelling of cross-sections (Figure 1C), we were able to identify fiber outlines and identify average cross-sectional area of all fibers using MyoVision 2.0. No significant differences were found between groups as the experiment was likely underpowered to find significant differences, although trends were clear when muscle cross-sectional area was plotted as a percent change between the left stimulated tibialis anterior muscle and right unstimulated contralateral control, (Figure 1F-G). This loss in muscle mass and fiber-cross-sectional area occurred without histological evidence of denervation, damage or degeneration as evidenced in Figure 1D by the absence of excess connective tissue, round fibers or central nuclei. We also observed little to no differences in the gene expression of *Myh3* and *Myh8* which encode embryonic and neonatal myosins respectively in mice and rats, Supplementary File 1. This suggests that there are similarities in whatever small damage response is generated following stimulation. NADH tetrazolium reductase staining, which is used as a proxy marker for mitochondrial content or oxidative capacity showed progressive increases with 2 (7.2 ± 1.6%), 10 (12.2 ± 5%) and 20 days (28 ± 9.1%) of daily training in the mouse as depicted in Figure 1E/H). These changes were similar in direction, but the magnitude of change was lower in the mouse compared to our previously published data using the same stimulation paradigm in rat.

**Figure 1:**
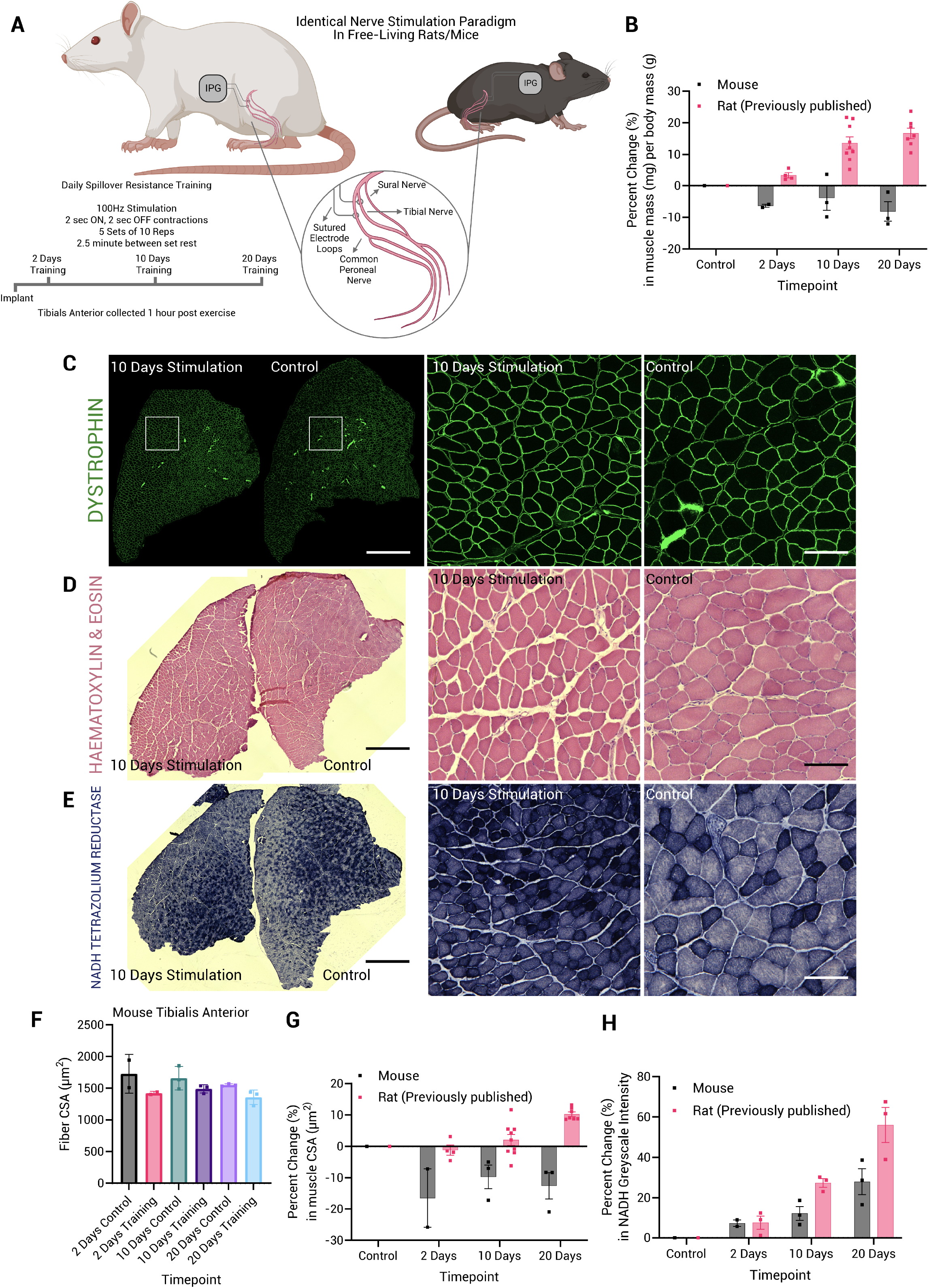
Experimental model and timecourse of changes in muscle mass with identical daily stimulation patterns in the rat and mouse. (A) Schematic of pulse generator surgical implantation and electrode placement. The cathode was placed underneath the common peroneal nerve, controlling motor activity in the dorsiflexors. The anode was placed underneath the tibial nerve, controlling motor activity in the plantarflexors. The ‘SpillOver’ stimulation paradigm allows for partial activation of the larger plantarflexor muscle group, providing a resisted contraction in the dorsiflexors. (B) Percent changes in muscle mass (mg) per body mass (g) over time in response to daily training between stimulated limbs and unstimulated contralateral controls (Mean ± Standard Deviation). (C) Representative dystrophin immunofluorescent labelling of a 10 day stimulated mouse muscle and unstimulated contralateral control. (D) Representative hematoxylin and eosin stain from 10 day stimulated muscle and unstimulated contralateral control muscle. (E) Representative NADH tetrazolium reductase staining from 10 day stimulated muscle and unstimulated contralateral control muscle. Scale bars indicate 2000µm/2mm for whole cross-sections and 100µm for regions of interest. (F) Fiber cross-sectional area µm^2^ measurements from whole cross-sections of mouse tibialis anterior assessed automatically using MyoVision 2.0. (G) Percent changes in muscle cross-sectional area in µm^2^ between unstimulated muscles in the right leg and stimulated muscle in the left leg. (H) Percent changes in NADH greyscale intensity between unstimulated muscles in the right leg and stimulated muscle in the left leg.

### Timecourse assessment of transcriptional responses to identical daily stimulation patterns in the rat and mouse

To understand the potentially divergent mechanisms resulting in substantial growth in stimulated rat muscle, but lack of hypertrophy in the mouse, we performed RNA-sequencing on the TA muscles from the mice. These responses were compared to an existing dataset in the rat, but the rat data was newly realigned to the latest rat genome assembly (Rn7). Rn7 has significant advantages over the previous Rn6 release of the rat genome as previously discussed^30,31^, which justify a new investigation of the rat dataset and allow a more appropriate comparison with transcripts of the mouse genome.

Our principal components analysis showed a clear distinction in both mice and rats between control and the acute exercise response at different levels of training status (Figure 2A/D). Using a 5% FDR cut off to identify significant DEGs versus the group of contralateral control muscles, we identified 3095, 3081 and 2293 genes in the acute response to exercise after 2, 10 and 20 days of training respectively in the rat, (Figure 2B). The number of overlapping genes between the 2- and 20-day response was 1023 genes, indicating that the acute exercise response changes significantly during a period of daily training in the rat. This is further highlighted in Figure 2C, where we show that only 1239 out of 4502 genes (27.5 %) were always differentially expressed at each timepoint, and thus independently of training status.

**Figure 2:**
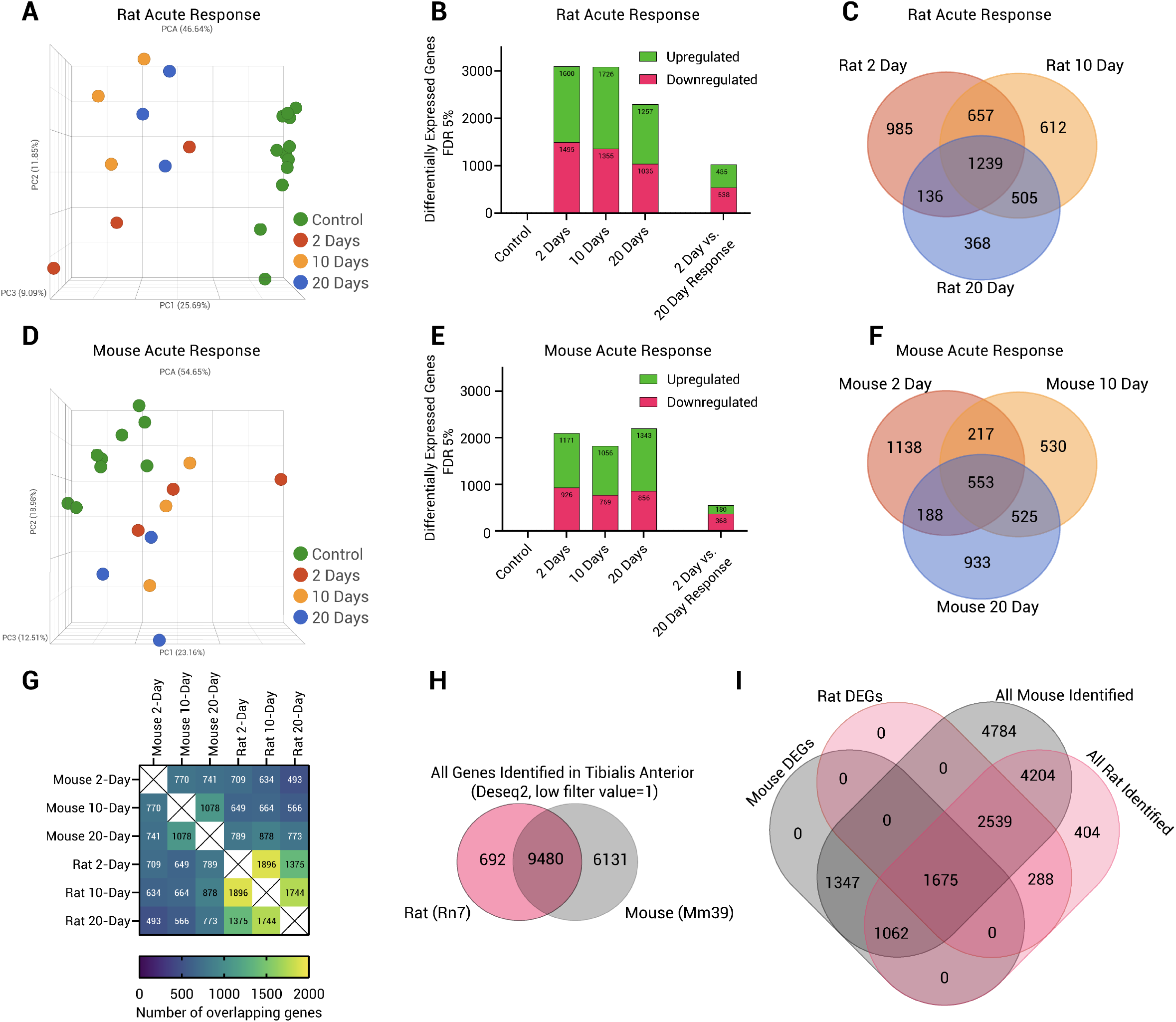
Summary of the transcriptional response across species in response to acute resistance exercise. (A) RNA-sequencing principal component analysis of the rat acute exercise response in relation to training status. (B) Differentially expressed genes relative to control in response to exercise in the rat using a false discovery rate of 5%. (C) Venn diagram analysis of training status specific responses to acute exercise in the rat. (D) RNA-sequencing principal component analysis of the mouse acute exercise response in relation to training status. (E) Differentially expressed genes relative to control in response to exercise in the mouse using a false discovery rate of 5%. (F) Venn diagram analysis of training status specific responses to acute exercise in the mouse. (G) Heatmap of differentially expressed genes overlapping between species, and stages of training. (H) Venn diagram analysis summarising the number of total genes identified from mouse (Mm39) and rat (Rn7) tibialis anterior muscles using the Deseq2 low filter value which removes genes with a mean low filter value of 1. (I) Venn diagram analysis overlapping the number of total differentially expressed genes in rat and mouse, as well as the number of genes identified in rat and mouse tissue. Data for figures 2C, F, H and I are available in supplemental file 2.

The total number of DEGs in the mouse in response to acute resistance exercise was about two thirds of the rat transcriptional response, even though the total number of genes identified in the transcriptome was greater for mouse than rat. Using the same statistical parameters as for the rat analysis to identify significant DEGs versus control muscles, we identified 2097, 1825 and 2199 genes in the acute response to exercise after 2, 10 and 20 days of training in the mouse, (Figure 2E). The number of overlapping genes between the 2- and 20-day response was 548 genes and only 553 genes (13.5 %) were consistently differentially expressed independently of training status (Figure 2F), showing the common phenomenon between species that the acute transcriptional response changed considerably as daily training progressed. Mouse DEGs and rat DEGs at each training status were compared and the number of overlapping features is presented in a heatmap (Figure 2G). A similar number of genes at each training status were found to be common between species, 709, 664 and 773 after 2, 10 and 20 days of training. Because there are known differences in genome annotation between species, we further probed our datasets to investigate whether the total number of genes expressed in the TA muscle (control or any stage of training) in the mouse and rat differed and how much of the basal transcriptome was conserved (using Deseq2 low filter value of <1). Interestingly, there were 692 genes only expressed in the rat TA (4.2 %), and 6131 specific to the mouse (37.6 %), but 9480 (58.2 %) which were expressed in both species, (Figure 2H). Overall, 15,611 genes were identified in the mouse using this filter and 10,172 in the rat.

To investigate whether the differences in the total number of identified genes influenced the number of DEGs between species we performed a Venn diagram analysis of the total number of mouse DEGs, the total number of rat DEGs and the total number of identified genes in mouse and rat (Figure 2I). Of the total number of genes identified (16303), 404 (2.5 %) were only identified in the rat but not differentially expressed with training, 4784 (29.3 %) were only identified in the mouse but not differentially expressed with training and 4204 (25.8 %) were expressed in both species but not differentially expressed in response to acute resistance exercise in one or both species. 288 genes (1.8 %) were only identified in the rat and were differentially expressed. Conversely, 1347 (8.3 %) of genes were only identified in the mouse and were also differentially expressed. Investigation of these genes revealed that most of them had no common ortholog in the other species, the gene had uncertain function, or they had uncharacterised locations within the genome assembly. A larger percentage of these were as expected found only in the mouse, suggesting that the many rat orthologs are yet to be identified in the genome assembly. For these reasons we chose genes identified in tibialis anterior muscles of both species for further investigation. Of genes identified in both species, 1062 (6.5 %) were only differentially expressed in the mouse, 2539 (15.6 %) were only differentially expressed in the rat and 1675 (10.3 %) were differentially expressed in both species.

### Common training status-specific transcriptional responses to resistance exercise in the rat and mouse

To identify potential differences that might explain the different growth responses between species, we overlapped the DEGs at each training time point between species to identify common and species-specific elements changing in the transcriptome. In the 2-day acute exercise response, only 709 (16.1 %) of the total number of DEGs in rats and mice were found to be common between species (Figure 3A). Pearson’s correlation of the Log^2^ fold changes in the common DEGs revealed a significant positive correlation between species (R = 0.71, *P* <0.0001), (Figure 3B). Analysis of these common and largely correlated features revealed enrichment of pathways related to stress responses, AMPK signalling, immune responses, autophagy, protein processing in the endoplasmic reticulum, glycogen metabolism, protein metabolism, and various processes involved directly or upstream of ribosome regulation and FoxO signalling pathways, (Figure 3C).

**Figure 3:**
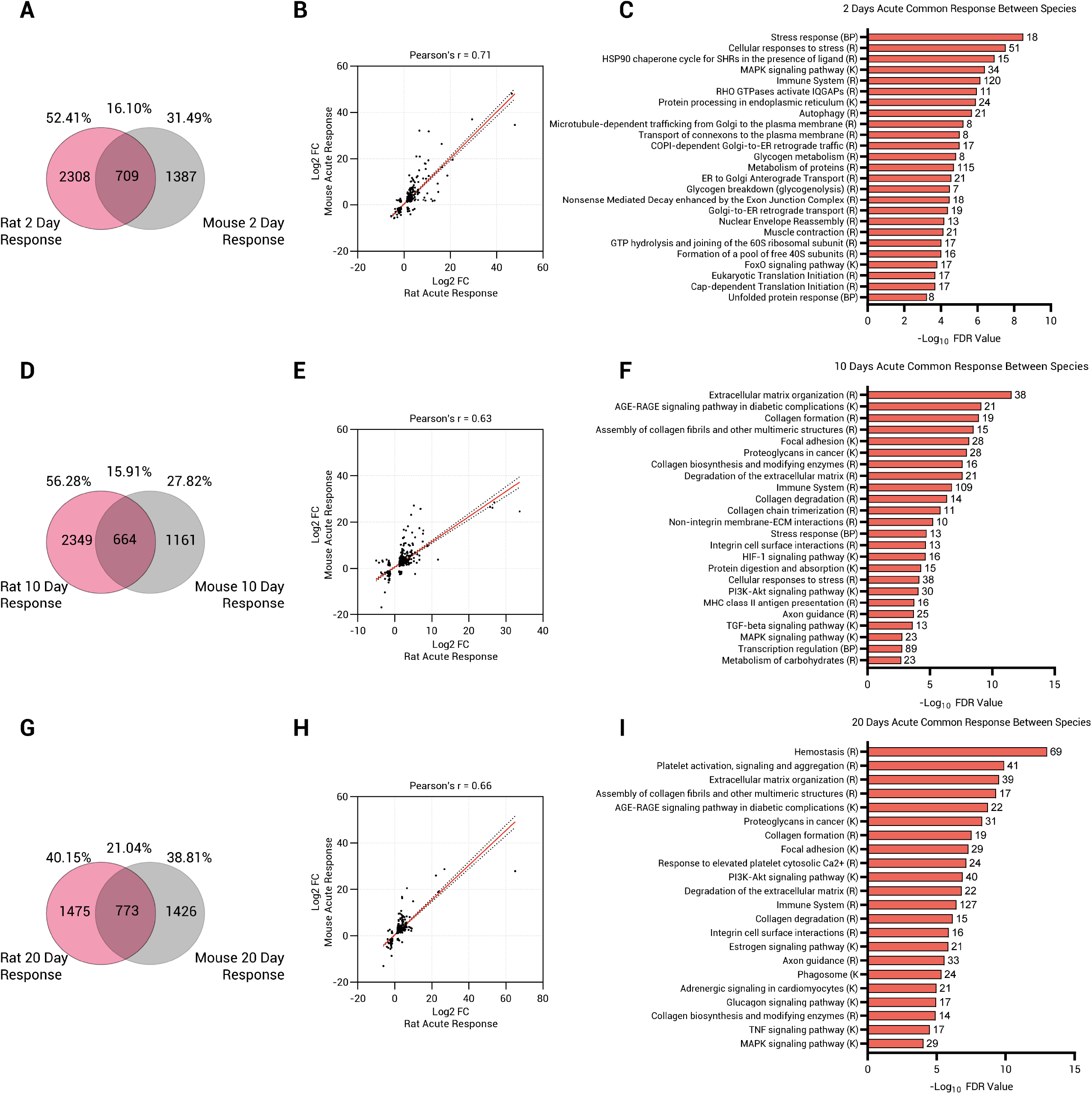
Common training status specific acute transcriptional responses to resistance exercise in rats and mice. (A) Venn diagram analysis of differentially expressed genes (5% false discovery rate) in response to acute exercise between mouse and rat after 2-days of training. (B) Pearsons correlation (Log^2^ FC normalised) of the 2-day acute exercise response of the shared differentially expressed genes identified in the figure 3A Venn diagram. (C) Pathway analysis of the shared features identified in figure 3A. (D) Venn diagram analysis of differentially expressed genes (5% false discovery rate) in response to acute exercise in mouse and rat after 10-days of training. (E) Pearsons correlation (Log^2^ FC normalised) of the 10-day acute exercise response of the shared differentially expressed genes identified in the figure 3D Venn diagram. (F) Pathway analysis of the shared features identified in figure 3D. (G) Venn diagram analysis of differentially expressed genes (5% false discovery rate) in response to acute exercise between mouse and rat after 20-days of training. (H) Pearsons correlation (Log^2^ FC normalised) of the 20-day acute exercise response of the shared differentially expressed genes identified in the figure 3G venn diagram. (I) Pathway analysis of the shared features identified in figure 3G. Pathways and their associated databases are annotated as follows KEGG Pathway (K), Reactome Pathway (R), GO Biological Process (BP). The number of genes associated with each pathway is presented adjacent to each pathway. Data for figures 3A-I are available in supplemental file 2.

In the 10-day acute exercise response, 664 genes (15.9 %) were commonly differentially expressed between species (Figure 3D), and they showed a similar significant correlation (R = 0.63, *P* <0.0001), (Figure 3E). However, pathway analysis revealed enrichment of a different set of pathways than in the 2- day acute exercise response. These included processes related to the extracellular matrix and collagen regulation, focal adhesion, proteoglycans and PI3k-Akt signalling pathways (Figure 3F).

Again, the 20-day acute exercise response showed a similar percentage of DEGs common to both species (773, 21 %) as previously noted with the 2- and 10-day responses. A similar significant positive correlation was identified in this gene set between species (R = 0.66, *P* <0.0001). A number of enriched pathways shared similarities to the 2- and 10-day responses, but hemostasis, platelet activation, signaling and aggregation pathways were specifically enriched at the 20-day timepoint. Many of the genes contained in these pathways are closely related to extracellular matrix and integrin remodeling.

We next investigated the genes involved in each identified pathway that showed common responses between species, plotting their Log^2^ Z-scores in response to acute exercise across the training timecourse. Genes involved in the stress response, including genes classified as immediate-early genes that are transiently activated by a wide variety of cellular stimuli, showed strikingly concordant changes in expression in response to acute resistance exercise across this timecourse and thus with similar training status, (Figure 4). Most of these genes are increased to their highest level in the 2-day exercise response with a gradual decline as training status increases. This pathway includes a number of transcription factors (*Ankrd1, Atf3, Csrnp1, Egr1, Fos, Hif1a, Jun, Junb, Jund, Myc, Nfatc3*), transcription co-activators (*Ppargc1a*), insulin-Akt-AMPK regulators (*Irs2, Mapk9*), nuclear receptors (*Nr4a1, Nr4a2, Nr4a3*) and heat shock factors (*Hspa1a, Hspa1b, Hspb1, Hspb2, Hspb6, Hspb7, Hspb8*) which are known to regulate a wide array of biological processes including metabolism and growth, (Figure 4A).

**Figure 4:**
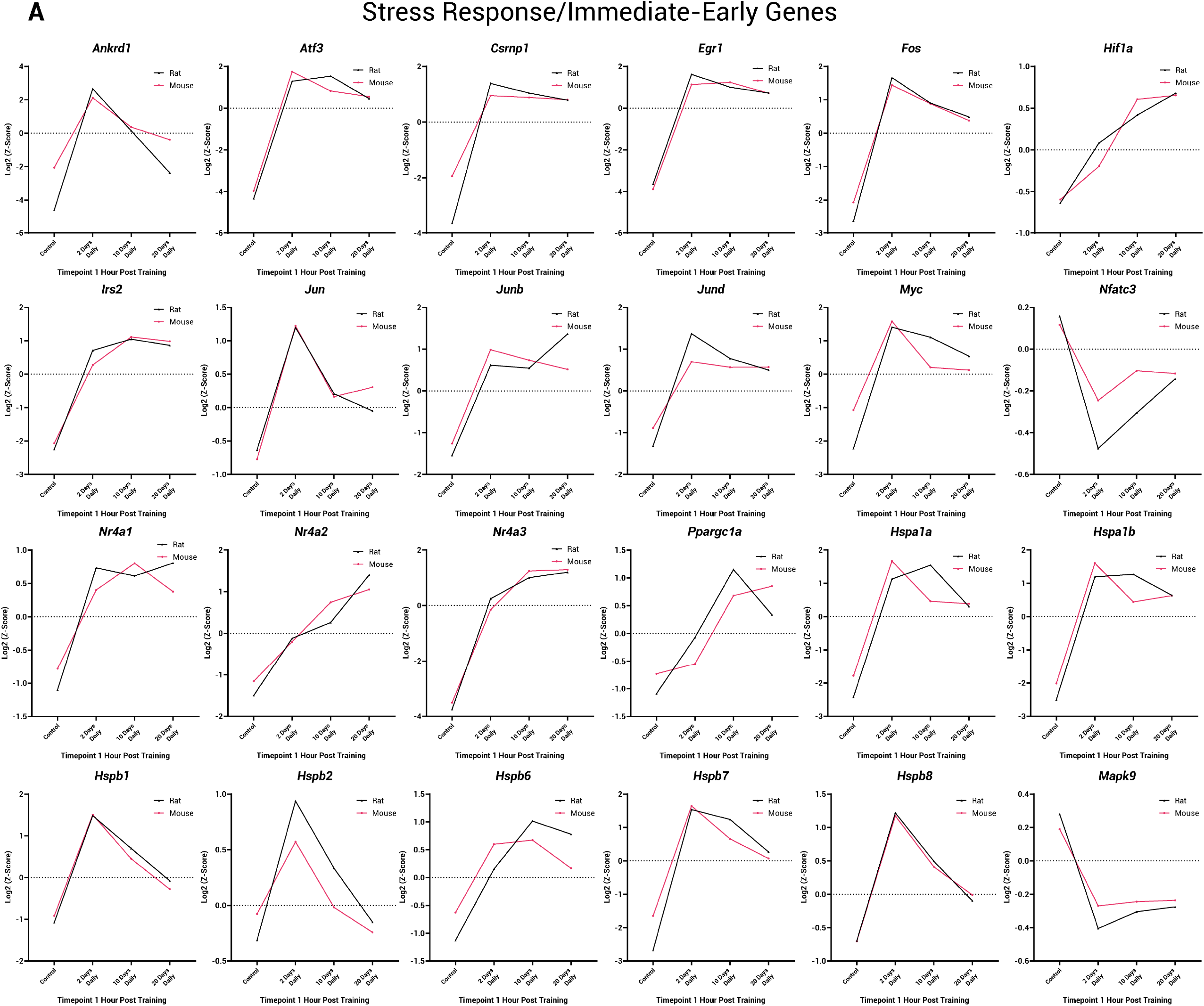
The timecourses for expression of stress response and ‘immediately early genes’ show strong similarities between species in response to resistance exercise. (A) Log^2^ z-scores of the Deseq2 normalised transcript abundance in control muscle and muscle after acute exercise with 2, 10 and 20 days of training history. Rat gene responses are indicated by a black line, mouse gene expression levels are indicated by pink lines.

Similarly, a large number of collagen and extracellular matrix associated genes show notable concordance between species and with training status including genes encoding enzymes involved in collagen cleavage, synthesis and maturation (*Adamsts2*, *Ctss*, *Loxl1*, *P4hb*, *Plod2*, *Serpinh1*), as well as collagens themselves (*Col1a1, Col1a2, Col3a1, Col4a1, Col5a1, Col5a2, Col6a2, Col8a1, Col11a1, Col14a1 Col15a1, Col18a1*), (Figure 5A). The majority of the genes within these pathways were most responsive to acute exercise between 10 and 20 days of training.

**Figure 5:**
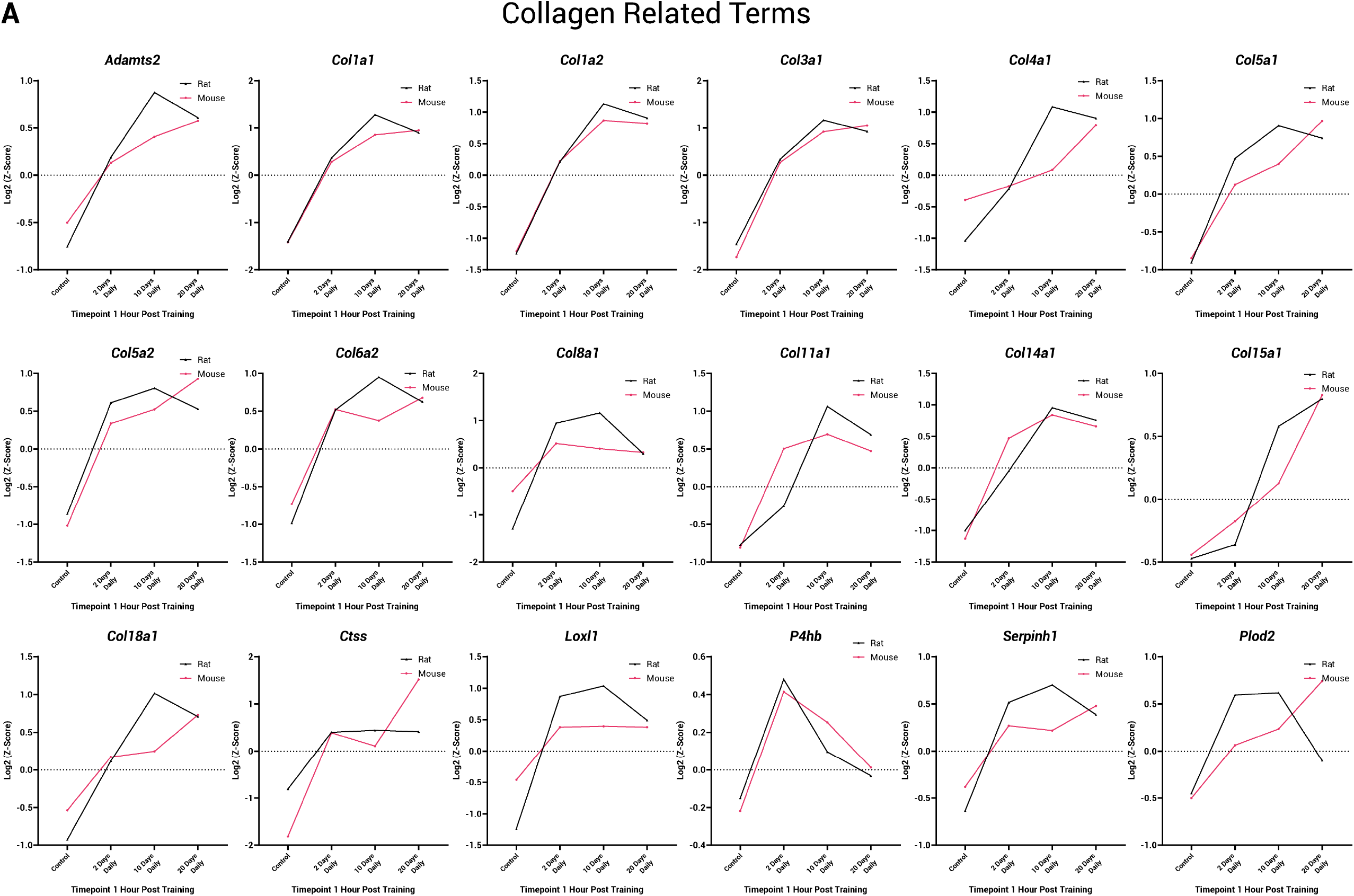
Collagen and extracellular matrix related genes show strong similarities between species and are progressively upregulated in response to resistance exercise. (A) Log^2^ z-scores of the Deseq2 normalised transcript abundance in control muscle and muscle after acute exercise with 2, 10 and 20 days of training history. Rat gene expression is indicated by black lines, mouse gene expression is indicated by pink lines.

A number of other pathways showed remarkably similar responses to acute exercise between species including genes associated with translation initiation and large and small ribosomal subunits (*Eif4a1, Rpl3, Rp4, Rpl12, Rpl13a, Rpl15, Rpl18, Rpl23, Rpl34, Rplp0, Rps12, Rps15, Rps18, Rps20, Rpsa, Rps26*), (Figure 6A). These associated genes showed patterns of expression similar to the stress response genes, peaking after just 2 days of stimulation before gradually declining (Figure 4). These early increases in ribosomal gene expression are a common feature of muscle hypertrophy, as ribosome abundance is a primary determinant of increased protein synthesis and translational capacity. We see them enchanced in both rat and mouse, but without muscle growth in the mouse. Similarly, 23 genes (*Capn2, Ccnd1, Diaph1, Flnc, Fn1, Ilk, Itga5, Itga7, Lamc1, Lamc2, Mapk9, Myl12a, Mylk2, Parvb, Pdgfb, Pxn, Src, Thbs1, Thbs3, Tln1, Vcl, Vegfa, Zyx)* associated with focal adhesion followed similar patterns in rat and mouse in response to acute exercise which suggests that the stimulation provides a similar mechanical stress in both species, (Figure 6B). Similar strong concordance was found in genes associated with autophagy, (Figure 6C). This gene set included a number of microtubule associated genes (*Dynll1, Map1lc3b, Tuba1a, Tuba1b, Tuba1c, Tuba4a, Tubb2b, Tubb4b, Tubb6, Vim)*, and genes associated with mitochondral control/mitophagy (*Pink1, Prkn, Ulk1, Vdac1*). Interestingly *Prkn,* an E3 ligase involved in removal of damaged mitochondria, was the only gene within this group to show discordance being significantly upregulated in the mouse after 2-days but signficantly downregulated in the rat before converging to similar levels after 10 and 20 days of exercise. Further similarities between species were observed for pathways broadly associated with ‘glucose/glycogen metabolism’ including 10 genes encoding important proteins involved in glucose transport, glycogenolysis and glycogen synthesis (*Agl, Calm1, Gbe1, Hk2, Pgm1, Phka1, Phkb, Phkg1, Pygm, Tbc1d1*) and 12 genes associated with ‘TGF-beta signalling’, (*Acvr1b, E2f4, Fst, Id1, Id2, Ppp2ca, Rgma, Rpsk6kb1, Smad3, Tgfb1, Tgif1 and Thbs1*). Individual plots for these genes and genes in Figure 6A-C are presented in Supplemental Figures 1-5.

**Figure 6:**
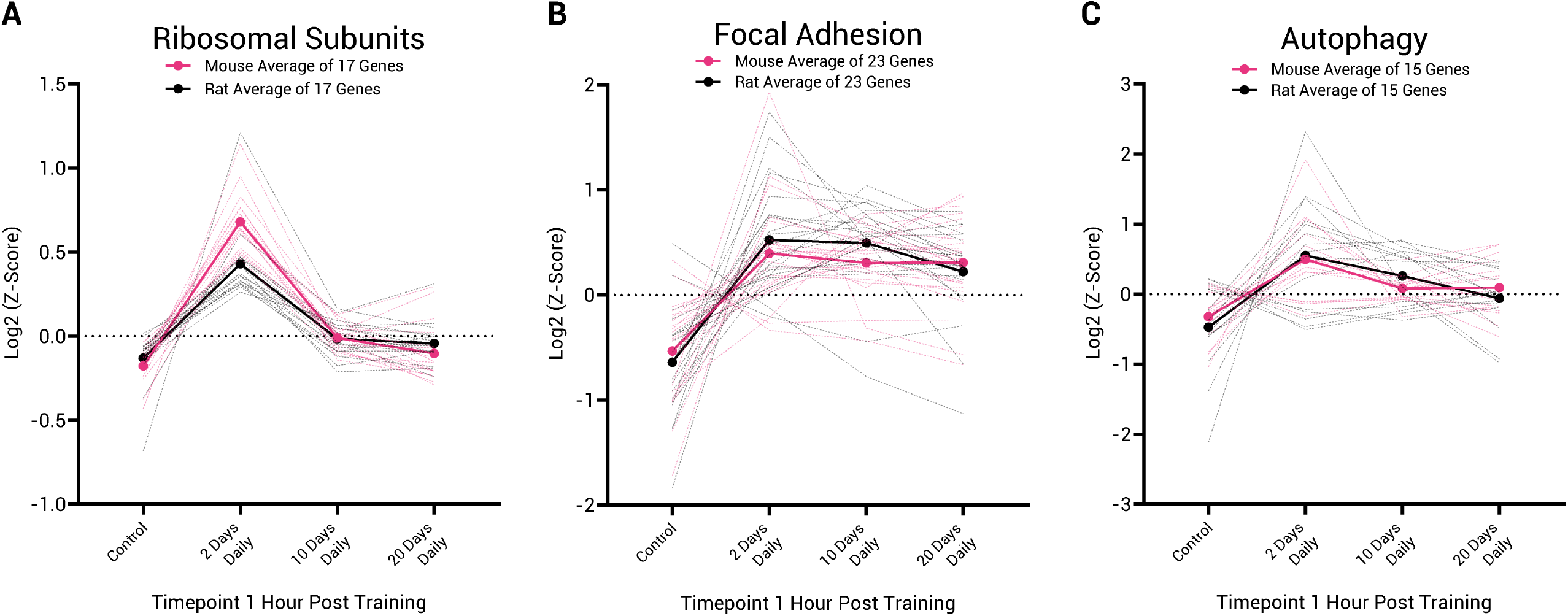
Strong concordance between species in gene expression related to ribosomal subunits, focal adhesion and autophagy in response to resistance exercise. Log^2^ z-scores of the Deseq2 normalised transcript abundance in control muscle and muscle after acute exercise with 2, 10 and 20 days of training. (A) Mean and individual gene traces for 17 genes associated with ‘Ribosomal Subunits’. (B) Mean and individual gene traces for 23 genes associated with ‘Focal Adhesion’. (C) Mean and individual gene traces for 15 genes associated with the term ‘Autophagy’. The mean Log^2^ z-score for the rat acute exercise responses is indicated by a solid black line, the mean Log^2^ z-score for the mouse acute exercise responses is indicated by a solid pink line. Individual rat gene responses are indicated by dashed black lines and individual mouse gene responses are indicated by dashed pink lines.

**Figure 7:**
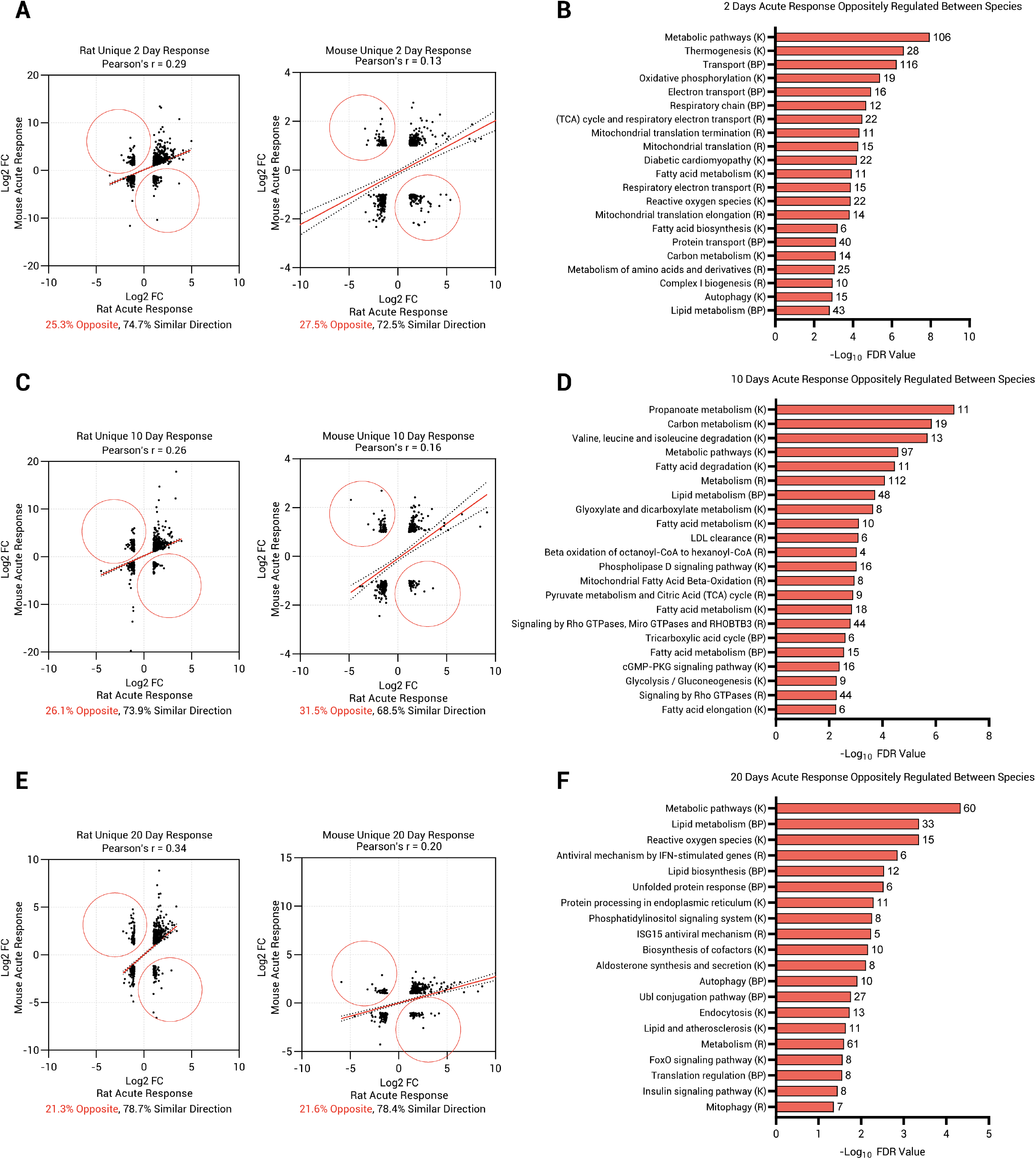
Species specific acute transcriptional responses to resistance exercise in rats and mice. (A) Genes identified to be differentially expressed (FDR = 5%) in response to acute exercise (2-day response) in only 1 species (Figure 3A) plotted as Pearsons correlations. (B) Pathway analysis of oppositely regulated genes (red circles) identified in the day 2 acute exercise response. (C) Genes identified to be differentially expressed (FDR = 5%) in response to acute exercise (10-day response) in only 1 species (Figure 3D) plotted as Pearsons correlations. (D) Pathway analysis of oppositely regulated genes (red circles) identified in the day 10 acute exercise response. (E) Genes identified to be differentially expressed (FDR = 5%) in response to acute exercise (20-day response) in only 1 species (Figure 3A) plotted as Pearsons correlations. (F) Pathway analysis of oppositely regulated genes (red circles) identified in the day 20 acute exercise response. Pathways and their associated databases are annotated as follows KEGG Pathway (K), Reactome Pathway (R), GO Biological Process (BP). The number of genes associated with each pathway is presented adjacent to each pathway. Data for figures 7A-F are available in supplemental file 2.

### Identification of divergent transcriptional responses between species to progressive resistance exercise

Although we identified a large number of genes showing similar responses to acute exercise in rat and mouse during a period of daily training, most DEGs were species specific in that they appeared as a DEG in one species but not the other (Figure 3A, D, G). However, we further filtered the species specific DEGs identified in this analysis by assessing Pearson’s correlations between the mouse and rat log^2^ fold changes in gene expression in response to acute exercise. Many genes had similar directionality of fold change in both species, despite not reaching the FDR cut off of 5% in one of the species. Across all training timepoints, between 68.5% and 78.7% of DEGs showed similar directionality, suggesting that a large proportion of the transcriptional response to acute exercise is conserved between mice and rats, yet it may differ in amplitude of response which results in missing an FDR cut-off of 5%. On the other hand, between 21.3 % and 31.5% of genes identified through this analysis were oppositely regulated between species, (indicated by red circles in Figure 5A, C, E).

In the 2-day acute exercise response (Figure 5A), 74.7 % of the rat 2-day DEGs showed similar directionality in the mouse and 72.5 % of the mouse 2-day DEGs showed similar directionality in the rat, despite not reaching significance in one species. 25.3 % and 27.5 % of rat and mouse DEGs respectively were oppositely regulated. In the 10-day acute exercise response (Figure 5C), 73.9 % of the rat 10-day DEGs showed similar directionality in the mouse and 68.5 % of the mouse 10-day DEGs showed similar directionality in the rat. 26.1 % and 31.5 % of rat and mouse DEGs respectively were oppositely regulated. Similarly in the 20-day acute exercise response (Figure 5E), 78.7 % of the rat 20-day response

DEGs showed similar directionality in the mouse, and 78.4 % of the 20-day mouse DEGs showed similar directionality in the rat. 21.3 % and 21.6 % showed opposing changes in gene expression in the 20-day acute exercise response comparison.

To examine further the biological processes represented by the oppositely regulated genes, we performed analysis to reveal enriched pathways, (Figure 5B, D, F). Interestingly, while there were transient changes in the enrichment of DEGs common to both species, the pathways enriched at 2, 10 and 20 days showed strong evidence for differences in metabolic processing in response to electrical stimulation between species. This included 106, 97 and 60 genes associated with ‘Metabolic Pathways’ after 2, 10 and 20 days of daily training respectively.13 of these genes were associated with ‘Carbon metabolism’, (*Acads, Acss2, Cat, Dlst, Esd, Gcsh, Gpt, Hibch, Mcee, Mdh1, Mdh2, Pdhb, Tkt*), (Figure 8A). A number of these genes are associated with the mitochondrion to provide substrate for the tricarboxylic acid cycle or to produce/convert substrates to tricarboxylic acid cycle intermediates - cleavage of glycine for degradation, production of pyruvate and glutamate, conversion of malate to oxaloacetate and facilitating the conversion of pyruvate to acetyl-coA. These components are upregulated in the mouse in the acute response to exercise after 2 and 10 days of training but are downregulated in the rat at the same timepoints suggesting that the metabolic demand of the identical activity pattern may be different in rat and mouse tibialis anterior muscle. Furthermore, we identified 21 genes associated with ‘Mitochondrial Translation’, *(Eif3k, Eif4a2, Eif5, Mrpl3, Mrpl4, Mrpl5, Mrpl6, Mrpl40, Mrpl41, Mrpl42, Mrpl44, Mrpl45, Mrpl51, Mrps5, Mrps14, Mrps15, Mrps21, Mtrf1l, Pabpc1, Rpl5, Srp72*), including genes involved in ribosome recruitment and translation initiation, large and small mitochondrial ribosomal protein complexes, and mitochondrial translation termination protein, (Figure 8B). Furthermore, as well as the protein synthetic machinery required to produce mitochondria, we also found discordance between species in 26 genes associated with ‘Oxidative Phosphorylation’, (*Atp5pb, Atp6v1a, Atp6v1e1, Cox5b, Cox7b, Dlst, Etfb, Gstz1, Hagh, Mdh2, Ndufa4, Ndufa8, Ndufb3, Ndufb4, Ndufb5, Ndufb6, Ndufc2,*

**Figure 8:**
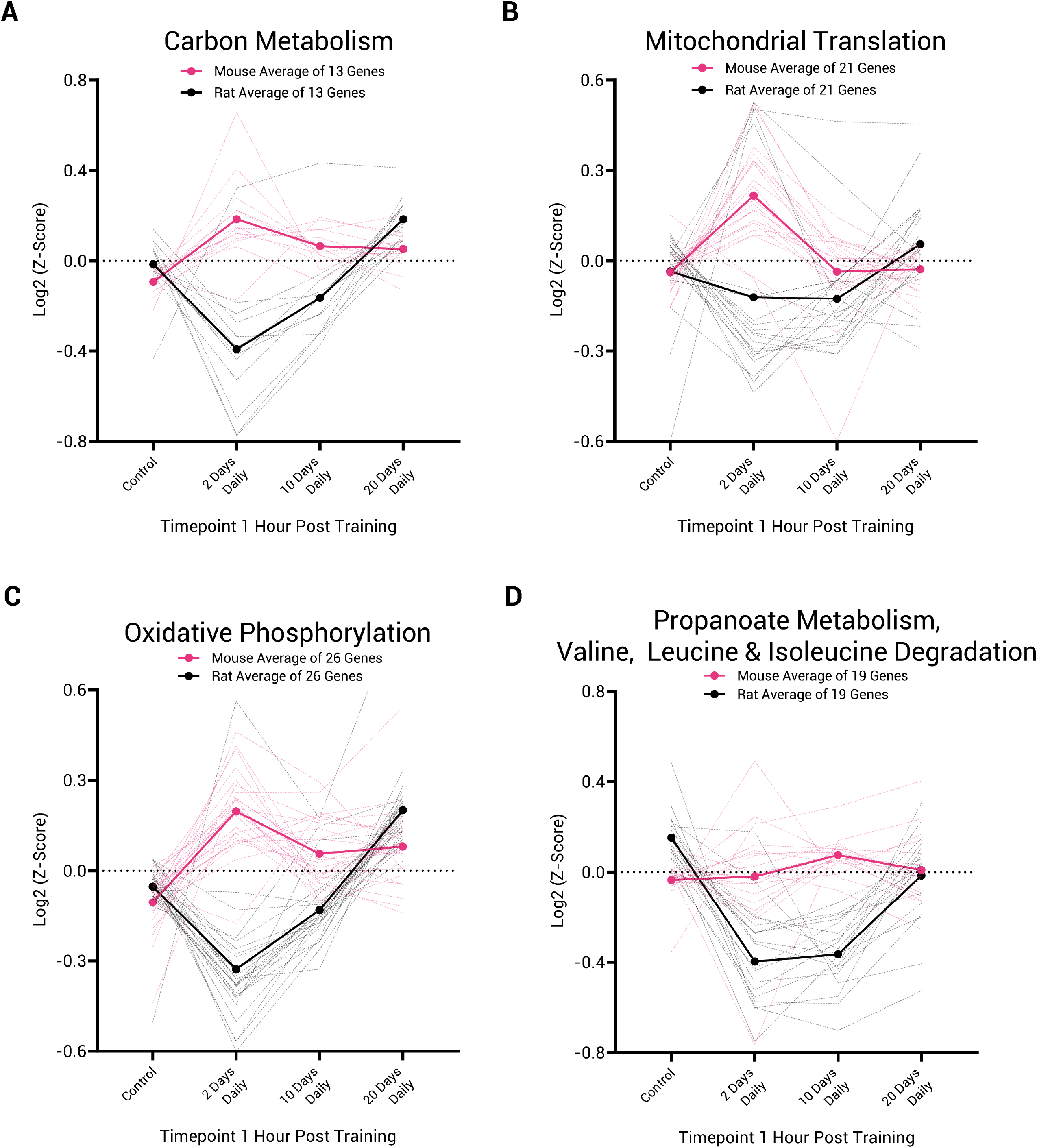
Metabolic pathways and mitochondrial translation are oppositely regulated between rat and mouse in response to acute resistance exercise. Log^2^ z-scores of the Deseq2 normalised transcript abundance in control muscle and muscle after acute exercise with 2, 10 and 20 days of training history. (A) Mean and individual gene traces for 13 genes associated with ‘Carbon Metabolism’. (B) Mean and individual gene traces for 21 genes associated with ‘Mitochondrial Translation’. (C) Mean and individual gene traces for 26 genes associated with the term ‘Oxidative Phosphorylation’. (D) Mean and individual gene traces for 19 genes associated with the term ‘Propanoate Metabolism, Valine, Leucine & Isoleucine Degradation’. The mean Log^2^ z-score for the rat acute exercise responses is indicated by a solid black line, the mean Log^2^ z-score for the mouse acute exercise responses is indicated by a solid pink line. Individual rat gene responses are indicated by dashed black lines and individual mouse gene responses are indicated by dashed pink lines.

*Ndufs3, Ndufs4, Ndufs6, Ndufv3, Pdhb, Ppa2, Sdhd, Surf1, Uqcr10*). This gene set included a number of mitochondrial complex associated genes, (Figure 8C). Again, this family of genes is upregulated in the mouse after 2-days of exercise, whereas it is consistently downregulated at all timepoints in the rat until 20-days when it is upregulated. As previously reported, the plateau in hypertrophy in the rat occurs after around 20-days when the muscle shifts to a more oxidative phenotype in response to Spillover resistance training. The observation that the mouse transcriptome seems to make this response immediately in response to resistance exercise training suggests that the mouse prioritises energy availability, in contrast to the rat which seems to prioritise muscle growth before a later shift to more oxidative metabolism. We can hypothesise that the exercise session produces a more profound energy deficit in the mouse TA than in the rat, and this hypothesis can be tested in future experiments.

A similar discordance between species was found in 19 genes associated with ‘Propanoate Metabolism and Valine, Leucine and Isoleucine Degradation’, (*Aacs, Acaa2, Acads, Acss1, Aldh2, Aldh9a1, Bckdha, Echdc1, Echs1, Hadh, Hadha, Hadhb, Hibadh, Ldha, Ldhb, Mccc1, Pccb, Sucla2, Suclg1*), (Figure 8D). A number of these genes are involved in regulation of energy metabolism and are again downregulated in the rat but remain either unchanged or partially upregulated in the mouse. Individual gene plots for figures 8A-D are available in supplemental figures 6-9. Further discordant pathways included ‘fatty acid/ lipid metabolism, degradation and biosynthesis’ ‘reactive oxygen species’, ‘phosphatidylinositol signalling’, ‘insulin related terms’, ‘foxo signalling’, as well as a number of pathways involved in ‘translation regulation’, ‘ubiquitin-like protein conjugation’ and the ‘unfolded protein response’. The genes included in these pathways are listed in Supplemental File 1.

Lastly, we chose to probe genes associated with muscle loss. Specifically we investigated genes commonly associated with the ubiquitin-proteasome system, genes associated with various forms of muscle atrophy and well-known markers of disuse caused by denervation or nerve silencing previously reported by our group and others^35–38^. Despite no histological evidence of denervation in the mouse, we sought to investigate any transcriptional changes that may infer that the muscle is programmed not to grow, despite a large number of anabolic gene signatures (Figure 3). While we recognise that changes in gene expression do not infer protein level or activity, we identified in the mouse tibialis anterior a large increase in the mRNA abundance of 11 of the 17 genes that encode alpha and beta proteasomal subunits of the 20s proteasome complex, all of which contribute to its assembly as a complex to remove protein substrates that are damaged or no longer needed, (Figure 9A). Most of these genes (*Psma1, Psma2, Psma4, Psma6, Psma7, Psmb1, Psmb3, Psmb4, Psmb5, Psmb5, Psmb6*), are unresponsive or downregulated in response to an acute training stimulus in the rat but are all substantially upregulated in the mouse in the 2-day acute response. Similar patterns are observed for the ATPase and Non-ATPase subunits of the 19s proteasome, and activator/inhibitor complexes, Figure (B-C). Genes encoding proteasome assembly chaperones are also upregulated in the mouse, but are seemingly unresponsive in the rat, Figure 9D. Again, while this does not infer proteasomal activity, transcribing genes costs energy so it is indicative of the need for greater degradation of proteins following exercise, or a greater need for 26s proteasome complexes to be available within the muscle fiber. The fact that this substantial increase is not found or is reduced in amplitude in the rat perhaps indicates how a robust hypertrophy is possible in response to electrical nerve stimulation.

**Figure 9:**
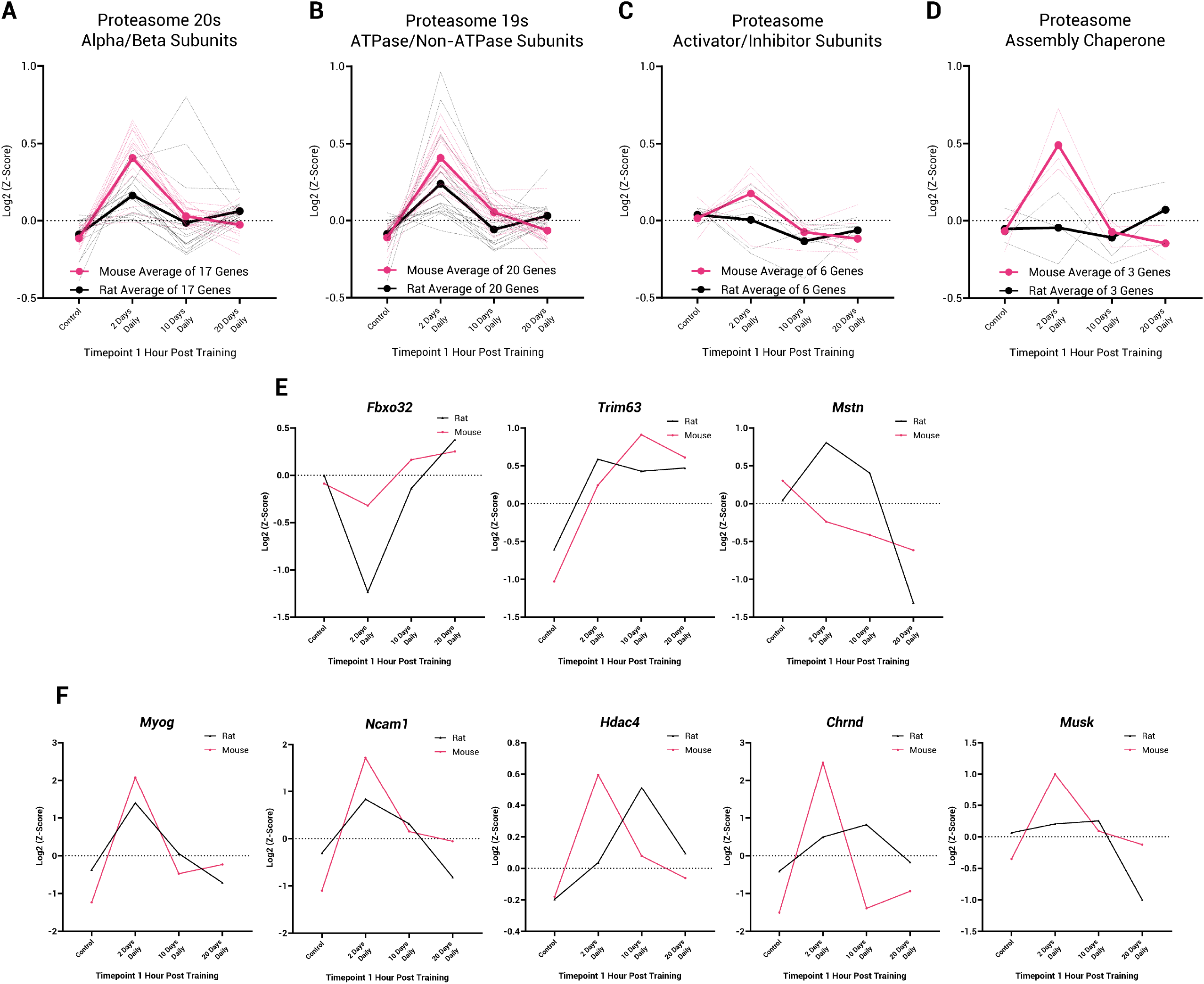
Greater increases in 26s proteasome and proteasome assembly chaperone gene expression in mice compared with rats may inhibit the early hypertrophic response. Log^2^ z-scores of the Deseq2 normalised transcript abundance in control muscle and muscle after acute exercise with 2, 10 and 20 days of training. (A) Mean and individual gene traces for 17 genes associated with ‘Alpha/Beta Subunits of the 20s proteasome. (B) Mean and individual gene traces for 20 genes associated with ‘ATPase and Non-ATPase Subunits of the 19s proteasome’. (C) Mean and individual gene traces for 6 genes associated with ‘Proteasome Activator/Inhibitor Complexes’. (D) Mean and individual gene traces for 3 genes associated with ‘Proteasome Assembly Chaperones’. The mean Log^2^ z-score for the rat acute exercise responses is indicated by a solid black line, the mean Log^2^ z-score for the mouse acute exercise responses is indicated by a solid pink line. Individual rat gene responses are indicated by dashed black lines and individual mouse gene responses are indicated by dashed pink lines. (E) Common Atrophy Associated Genes and (C) Genes associated with denervation. Rat gene expression is indicated by black lines, mouse gene expression is indicated by pink lines.

We also studied the timecourse of gene expression of a number of genes associated with muscle atrophy, denervation, and nerve silencing. As depicted in Figure 9E-F, we show that a number of these marker genes (*Mstn, Fbxo32, Trim63, Chrnd, Musk, Ncam1, Hdac4, Myog*) are upregulated in the mouse, and in this case, the same temporal patterns of change are observed in the rat. As the muscle grows in the rat, it is hard to postulate that this signature is indicative of denervation in the mouse when the same electrical stimulation pattern is provided in both. We note that the majority of these genes that have been reported to change with denervation are also acutely altered in the hours post exercise to regulate rates of protein synthesis and degradation. As our samples are taken 1-hour post exercise, further sampling of the basal transcriptome at each trained state prior to exercise will be important for future interpretations. Despite this, our histological analysis shows that the mouse muscles are healthy and the lack of growth following our training paradigm is most likely attributable to a net negative protein balance.

## Discussion

In this investigation, we identified concordance and discordance in the acute transcriptional responses between mice and rats to identical nerve stimulation parameters in free-living conditions. Skeletal muscle phenotyping over a timecourse of response to resistance exercise training revealed a progressive hypertrophy in the rat muscle, but the same stimulus had little effect on the mouse in terms of growth. We asked the question, did the mouse not mount a response to the exercise, or was the stimulus somehow ineffective in that species? No, in both species the stimulation generated visible contractions and the transcriptional response was complex and substantial with many common features but also striking differences. Combining the timecourse assessments with RNA-sequencing of the acute exercise response after 2, 10 and 20 training sessions, we found that there are large commonalities between the acute transcriptional response to exercise in mouse and rat tibialis anterior muscle across the timecourse of adaptation including stress responses, focal adhesion, collagen homeostasis, ribosomal biogenesis, and autophagy. However, we were able to identify species specific unique transcriptional responses mainly relating to metabolic processes, energy homeostasis and mitochondrial biogenesis, and these are therefore most probably related to the discordant growth response.

We identified remarkable concordance between both stress response and mechanical stress associated genes which is indicative that the contractile stimulus and the response to that stimulus are extremely similar between species. However, our data demonstrates that the appearance of some components of the transcriptional exercise response typically associated with muscle hypertrophy (immediate-early genes and ribosomal biogenesis), cannot be taken as sufficient evidence that subsequent hypertrophy will take place with repeated exercise of the same type, (Figures 3-6). Despite concordance in the mRNA level of key transcriptional regulators between species and with training status (*Ankrd1, Atf3, Csrnp1, Egr1, Fos, Hif1a, Jun, Junb, Jund, Myc, Nfatc3*, *Ppargc1a*, *Nr4a1, Nr4a2, Nr4a3*, *Hspa1a, Hspa1b, Hspb1, Hspb2, Hspb6, Hspb7, Hspb8*), further work should elucidate whether their activity is conserved between species. The strong concordance in collagen and collagen homeostasis related genes also suggests that the muscle undergoes similar structural remodelling in response to stimulation, yet a hypertrophic response is lacking.

We propose that one contribution to the lack of muscle hypertrophy in the mouse in response to programmed electrical stimulation occurs as a result of different metabolic requirements of the exercise bout which results in a different transcriptional response. We highlight that a major difference between these species is that although genes involved in glucose and glycogen metabolism, including transporters and rate limiting enzymes, show a concordant response in response to exercise, genes involved in carbon metabolism and leucine, isoleucine, and valine degradation are suppressed in the early response to acute exercise in the rat, but upregulated in the mouse. This may suggest that the mouse has an increased demand for anaplerotic substrates post exercise that are supplied through increased amino acid metabolism. This increased demand for free amino acids in muscle to replenish the citric acid cycle substrates may increase protein degradation in the mouse which precludes a growth response. We have not measured protein synthesis and degradation rates or net balance, but such measurements would prove informative in understanding the lack of hypertrophy. However, A number of papers have used similar paradigms, stimulating the entire sciatic nerve to elicit resisted high force contractions of the lower leg muscles^23,39^. These studies have shown that electrical stimulation can invoke an increase in protein synthesis in the hours following cessation of contraction. This has also been shown to be concomitant with the rapamycin-sensitive element of mTOR which signals to increase phosphorylation of several residues on P70 and S6 proteins^40^. Electrical stimulation is clearly capable of inducing a protein synthetic response and for growth this is often accompanied by a reduction in 20S proteasome activity. For example, following synergist ablation in mice, it has been recorded that 20S proteasome activity is reduced by ∼63% in the plantaris, and ∼20% in the soleus, resulting in a greater rate and magnitude of growth in the plantaris^41^. By contrast, our gene expression data infers an increased requirement for the 20S proteasome system (Figure 9) in the mouse after a resistance training bout. Again, we propose that the lack of growth following our training paradigm in the mouse is most likely attributable to a net negative protein balance.

Alongside the discordance in the genes associated with carbon metabolism and amino acid availability, the mouse transcriptome showed upregulation of a large network of genes involved in lipid oxidation, as well as genes encoding subunit constituents of all 5 mitochondrial complexes and proteins involved in their appropriate assembly. Genes encoding members of the pyruvate dehydrogenase complex, TCA cycle intermediate enzymes, oxidative metabolism and genes encoding proteins important for mitochondrial translation initiation and capacity were also upregulated. These are oppositely regulated in the rat, until 20-days of training when muscle hypertrophy has ceased. We did not directly test substrate availability, usage or metabolic enzyme activity/content after acute exercise in our model, but it is possible that the mouse tibialis anterior being less oxidative than the rat and rabbit tibialis anterior muscle^42^ promotes an upregulation of metabolic components even during or following our ‘typical’ resistance training protocol of 5 sets of 10 repetitions which may not be required to maintain homeostasis after stimulation with the same protocol in the rat.

Since our publication of the timecourse of change in the *acute* transcriptional responses to exercise over 4 weeks of daily training in the rat, there have been very few additional studies that investigate this important aspect of the exercise response. Furrer *et al.* reported from a study of endurance training in mice that while the basal (rested) transcriptomes of trained and untrained muscles are remarkably similar, training status substantially affects the acute transcriptional response to exercise^43^ including increases and decreases in the amplitude of gene expression, as well as changing the phase or rate of response post exercise (peaking earlier post exercise) as we have previously reported^17^. Training status was also reported to change the transcriptional program itself with the loss of response in some genes and production of de-novo responses of other genes as training status increases, similar to our report. To date, similar investigations into the modulation of transcription by training status have been limited to endurance or high intensity interval exercise responses^44,45^, and our new data and previous data from our group^17^ remain a valuable resource for understanding training status modulation of the acute transcriptional response to resistance-like exercise. Whether the mouse is a suitable model to study human muscle growth or wasting is an important and current research question. A recent transcriptomic survey of 4 muscles in mice and rats across the lifespan, suggests that rats mimic human aging and sarcopenia more closely than mouse^35^. Large differences in the progressive dysregulation of genes associated with metabolic processes and mitochondrial function were found in rat, but not in mouse suggesting that the use of the mouse or mouse tissue must be carefully interpreted when translating to aspects of human aging and sarcopenia. We propose that the lack of growth in the mouse in response to a daily session of electrical stimulation, which causes hypertrophy in rats and in humans^46–48^ suggests that work in the mouse needs careful interpretation when applied to human hypertrophy or for example, optimisation of neuromuscular electrical stimulation interventions for clinical use in humans.

We identify a number of limitations and future considerations within these experiments including the limited timepoints post exercise. We use 1-hour post exercise as it has previously been identified as the timepoint which yields the largest transcriptional changes in skeletal muscle^49^, but we cannot discount the possibility of different transcriptional responses occurring in different phases post exercise between species as recently reported^43^. We also understand that transcriptional regulation of mRNAs is one aspect of the adaptive response to exercise and accompanying protein and metabolomic data will help future understanding of what drives the transcriptional response and resultant changes in muscle size.

Overall, our findings illustrate that there are both concordant and discordant elements in the acute responses to exercise in the same muscle in two very closely related murine species. The exercise response is not uniform across species, despite using highly controlled, programmed exercise in free living animals, and differs with different stages of training. Further study of epigenetic control, transcription factor kinetics and post-transcriptional regulation of mRNAs will help to explain the hierarchy of the response illustrated in this study in which adaptation of the metabolic support for contraction in the mouse appears to be triggered before the net increase in protein generation that is required for muscle growth.

## Ethics declarations

This study received funding from DOD SCIRP IIR SC190031 (To CPC and JCJ), VA Rehabilitation Research and Development Service I50RX002020. The views represented in this manuscript are not reflective of the United States Government or the Department of Veterans Affairs.

## Data Availability

The data that support the findings of this study are openly available via the National Institutes of Health Gene Expression Omnibus. Raw RNA-sequencing data (FASTq) is available through GEO Accessions, GSE196147 and GSE228050. Processed, normalised count data for RNA-sequencing experiments is available in supplemental file 1. Any additional data are available upon request from the corresponding author.

## Supplemental Figure Legends

**Supplemental Figure 1:**
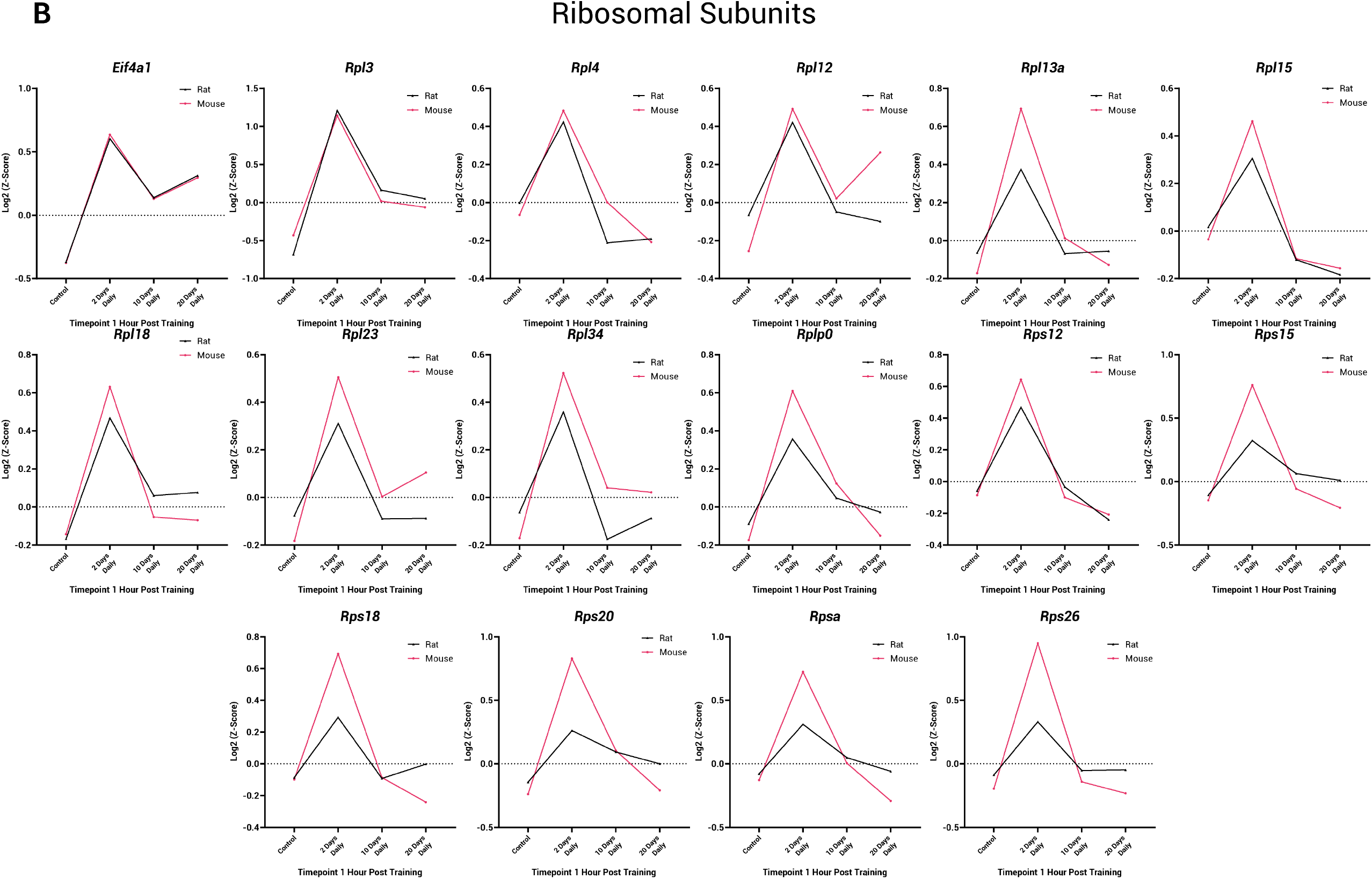
(A) Individual gene plots for genes associated with ribosomal subunits. Rat gene expression is indicated by black lines, mouse gene expression is indicated by pink lines. Genes are also presented overlayed in Figure 6.

**Supplemental Figure 2:**
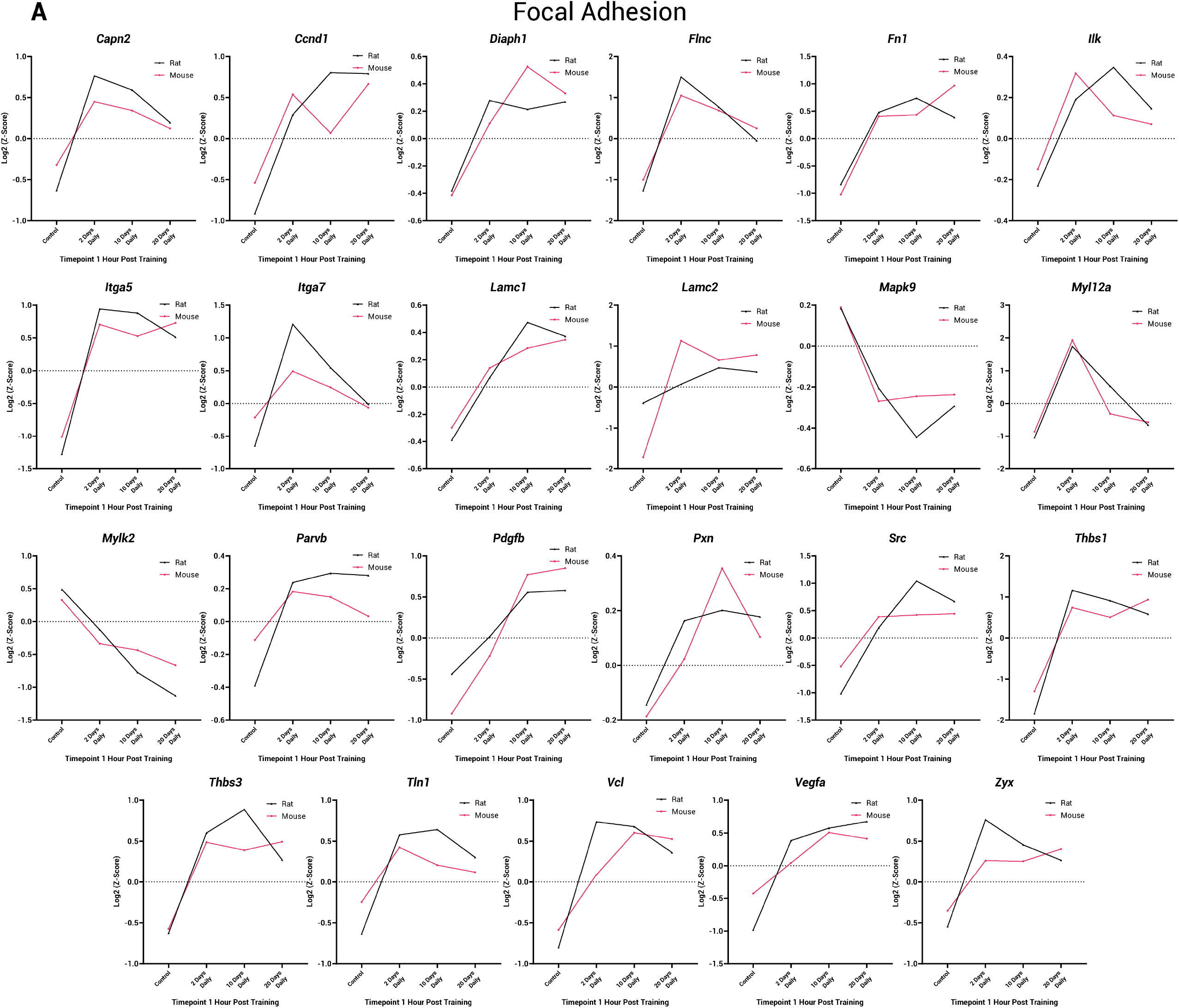
(A) Individual gene plots for genes associated with focal adhesion. Rat gene expression is indicated by black lines, mouse gene expression is indicated by pink lines. Genes are also presented overlayed in Figure 6.

**Supplemental Figure 3:**
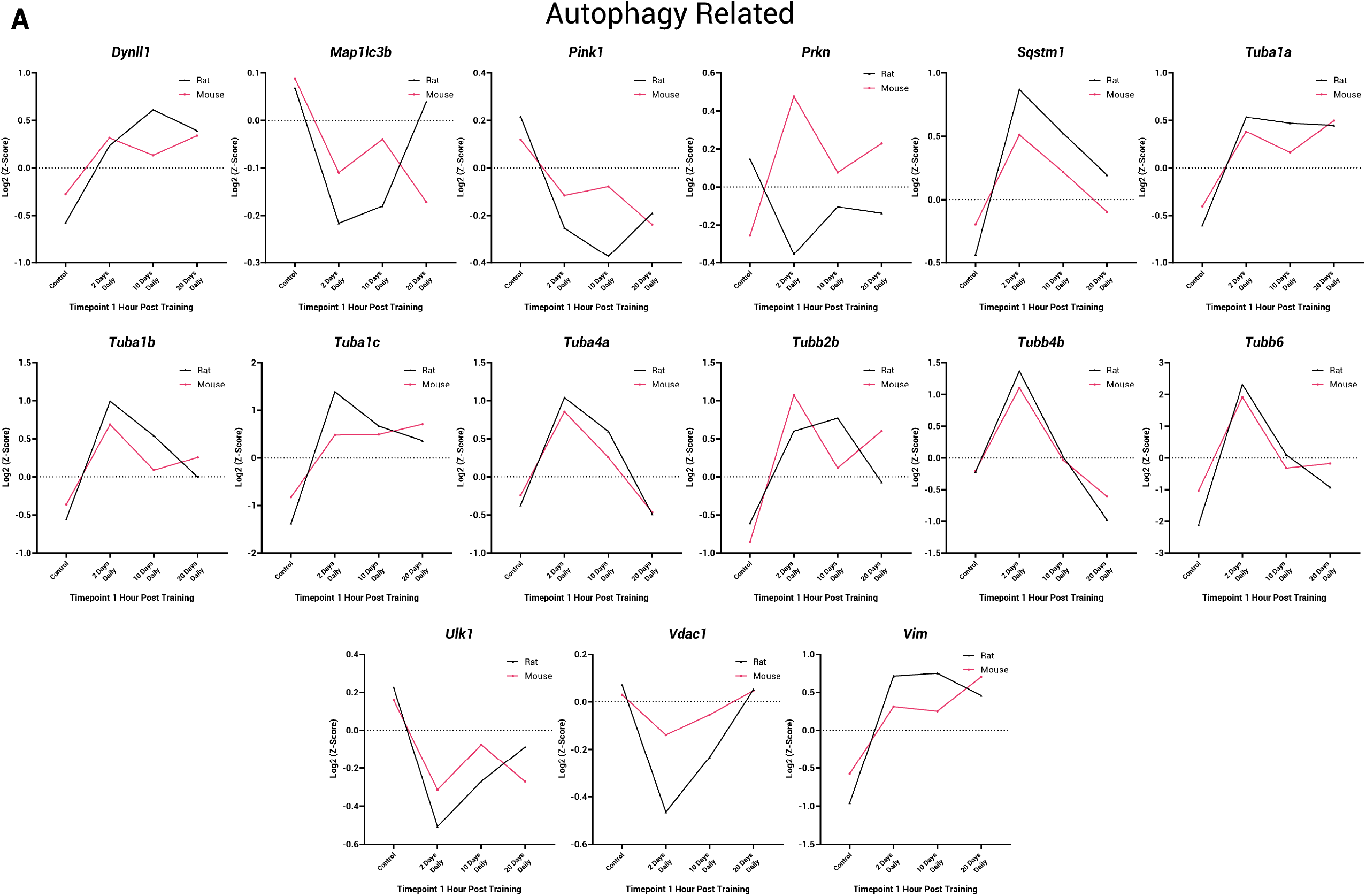
(A) Individual gene plots for genes associated with autophagy. Rat gene expression is indicated by black lines, mouse gene expression is indicated by pink lines. Genes are also presented overlayed in Figure 6.

**Supplemental Figure 4:**
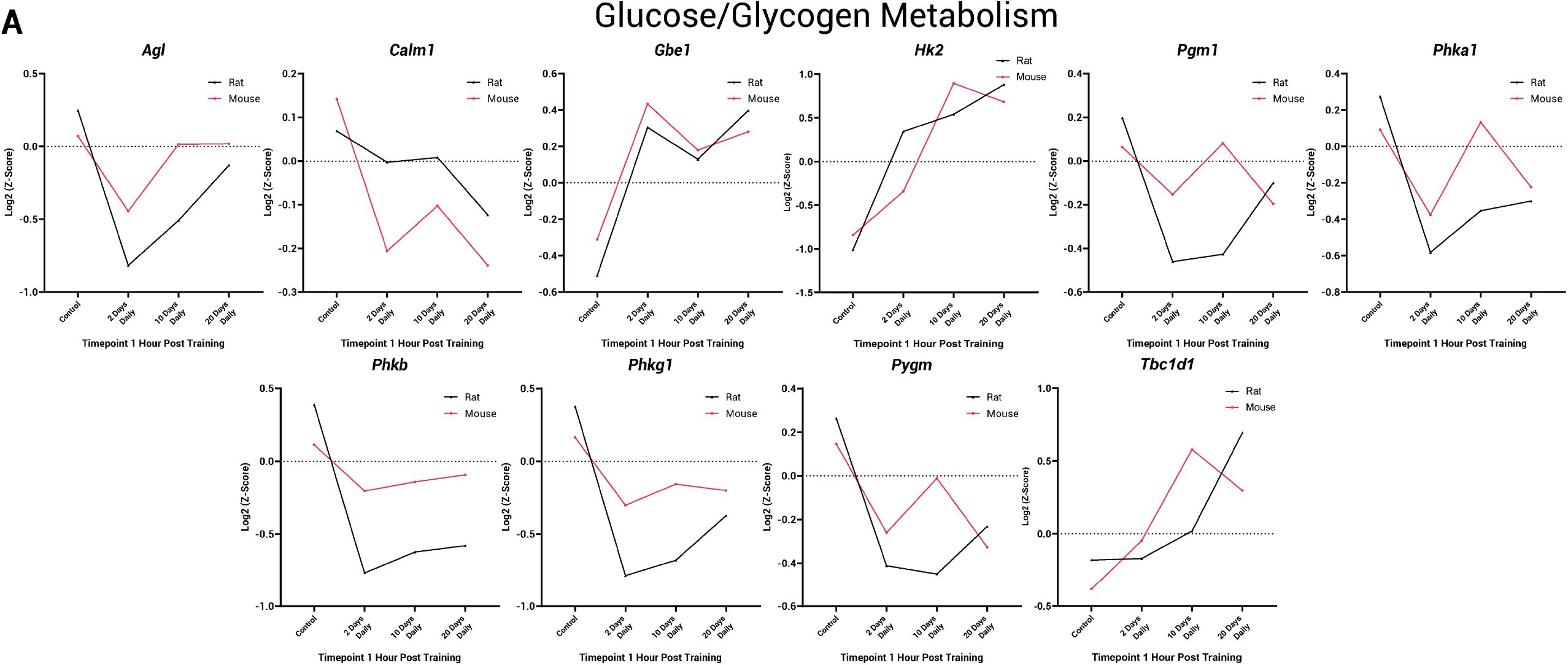
(A) Individual gene plots for genes associated with glucose and glycogen metabolism. Rat gene expression is indicated by black lines, mouse gene expression is indicated by pink lines.

**Supplemental Figure 5:**
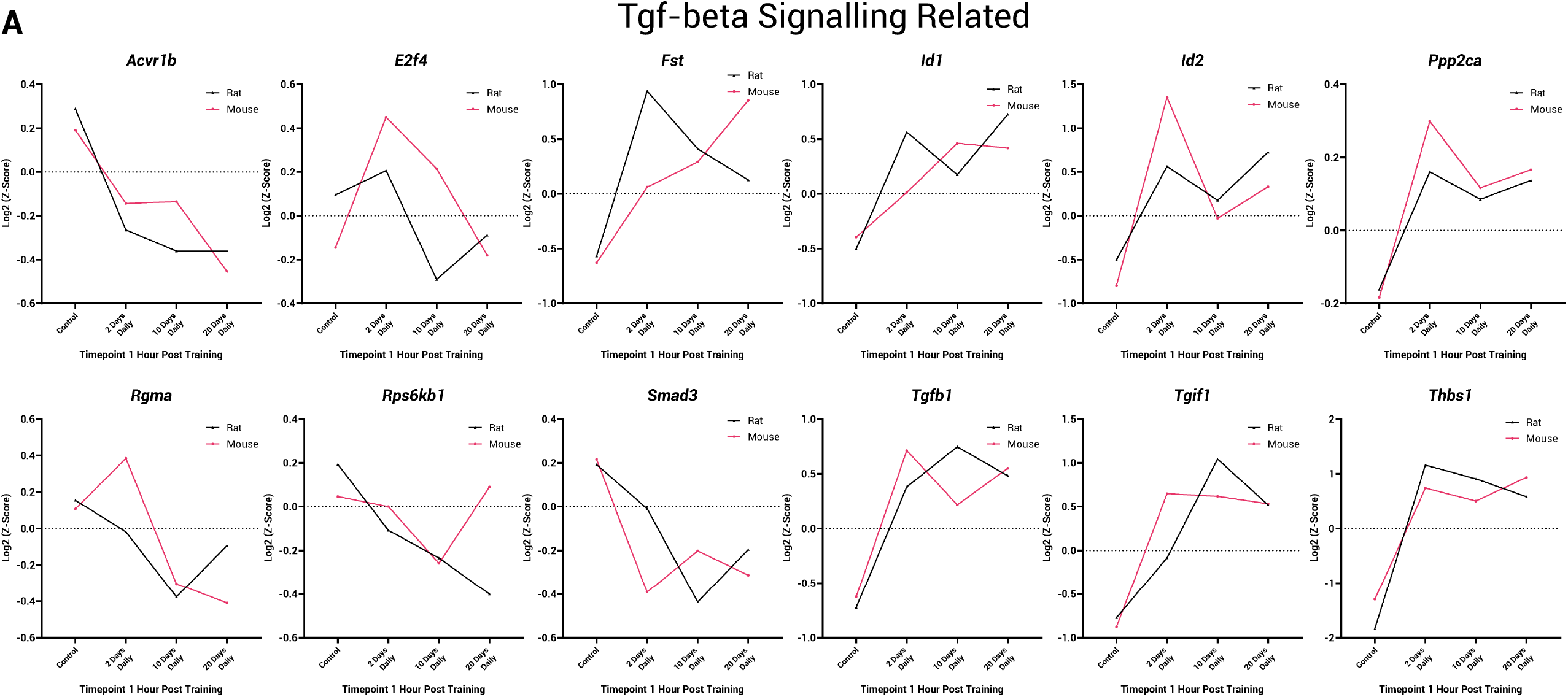
(A) Individual gene plots for genes associated with TGF-beta signaling. Rat gene expression is indicated by black lines, mouse gene expression is indicated by pink lines.

**Supplemental Figure 6:**
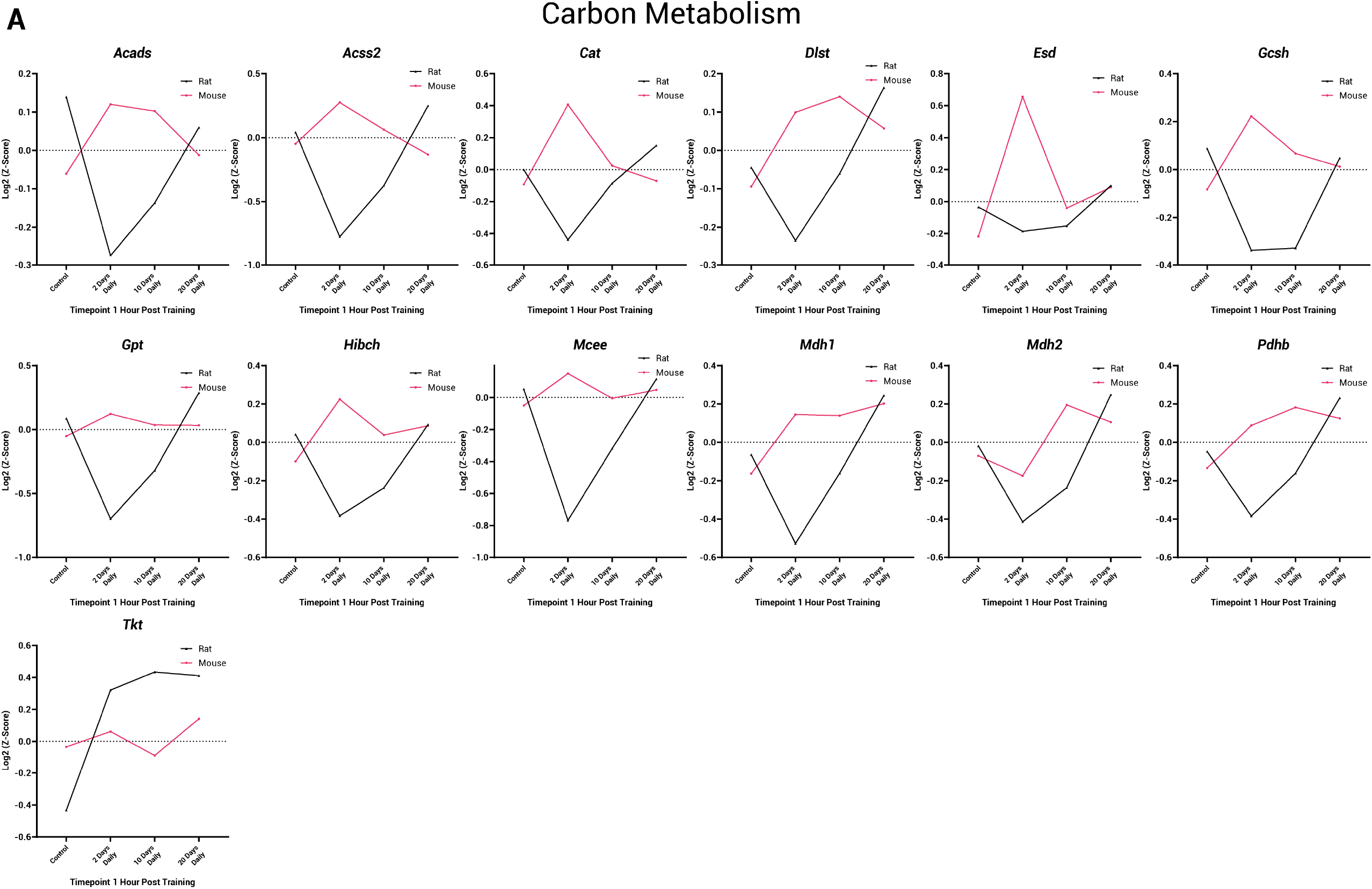
(A) Individual gene plots for genes associated with carbon metabolism. Rat gene expression is indicated by black lines, mouse gene expression is indicated by pink lines. Genes are also presented overlayed in Figure 8.

**Supplemental Figure 7:**
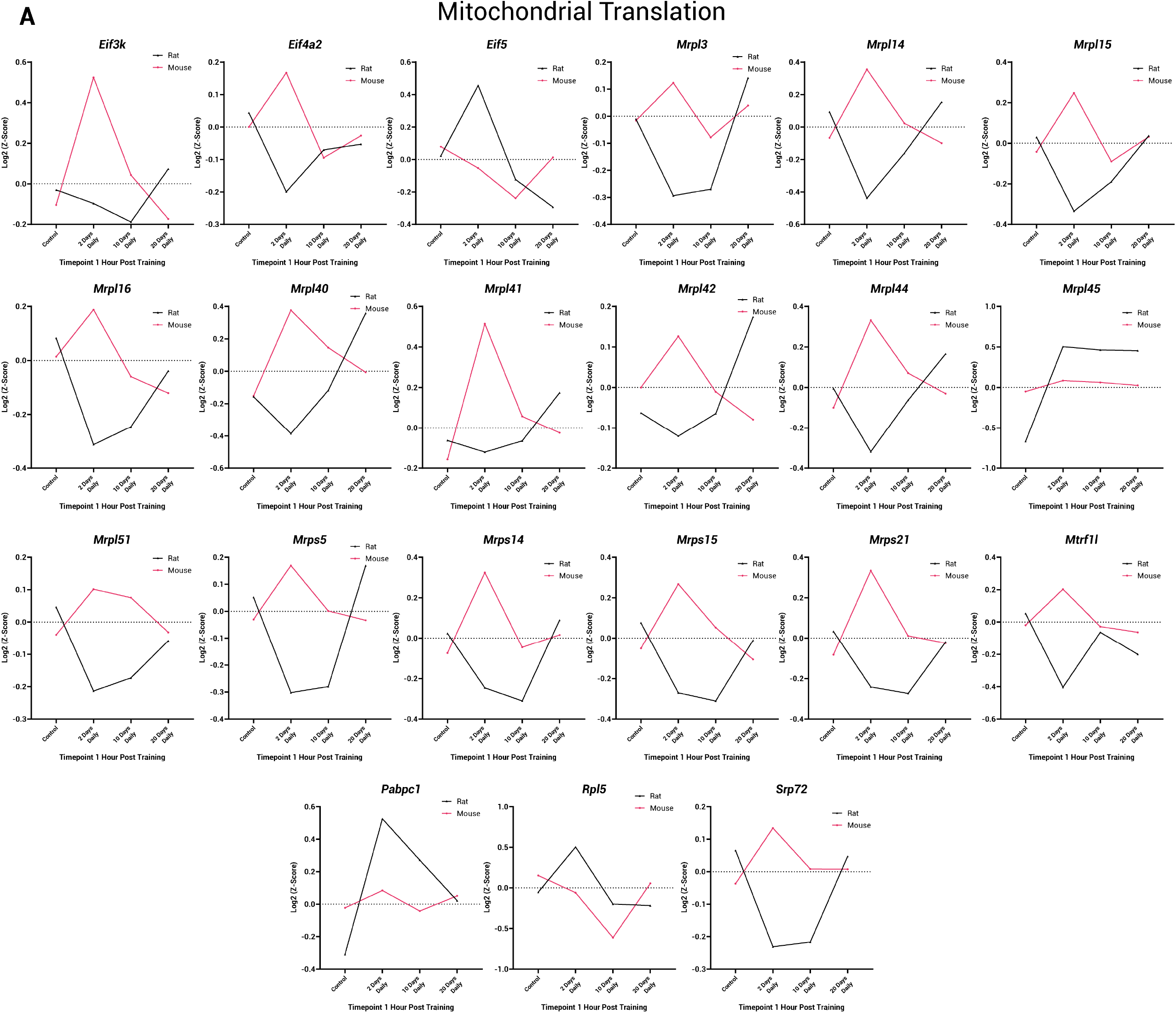
(A) Individual gene plots for genes associated with Mitochondrial Translation. Rat gene expression is indicated by black lines, mouse gene expression is indicated by pink lines. Genes are also presented overlayed in Figure 8.

**Supplemental Figure 8:**
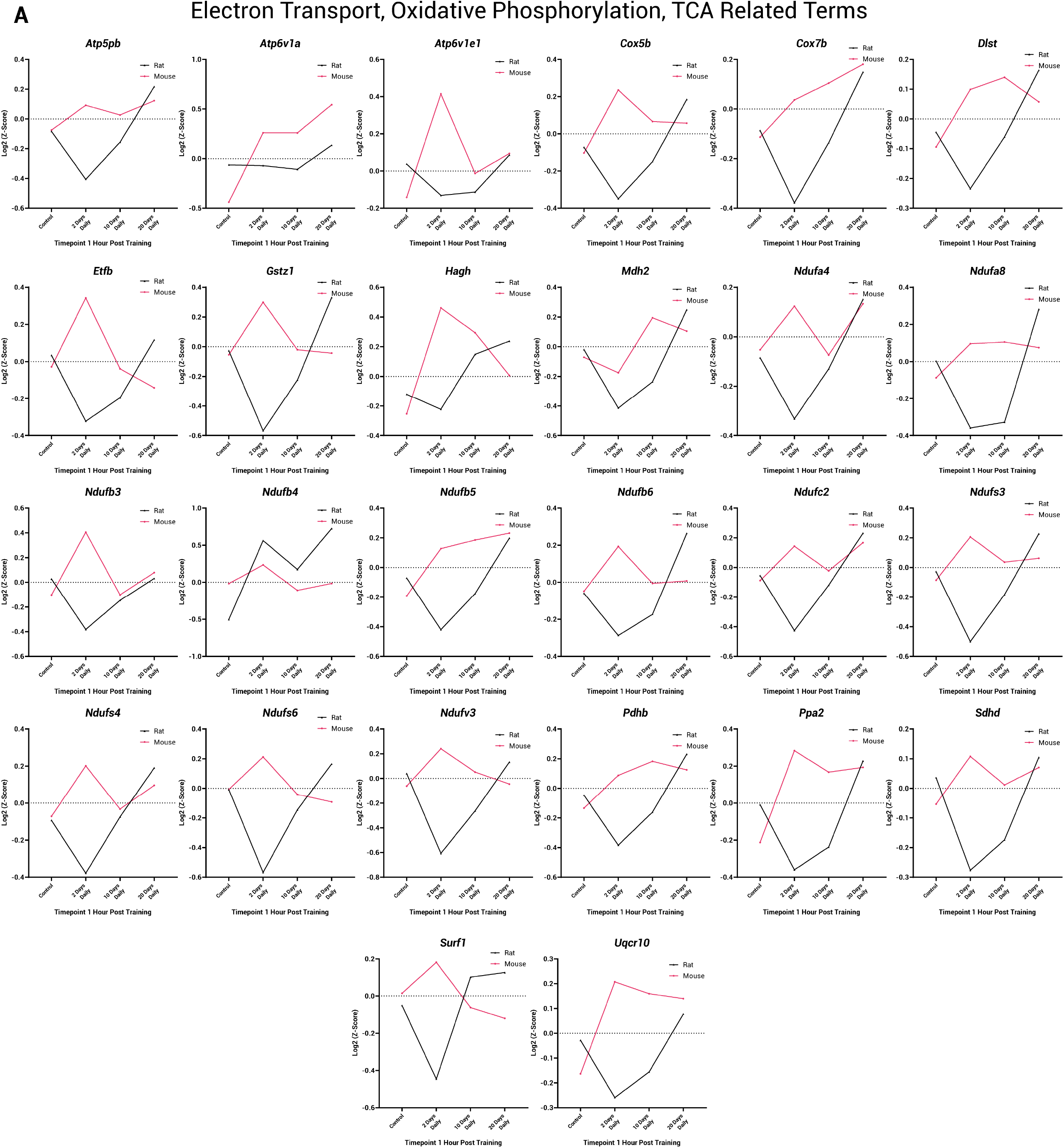
(A) Individual gene plots for genes associated with Electron Transport, Oxidative Phosphorylation and TCA related terms. Rat gene expression is indicated by black lines, mouse gene expression is indicated by pink lines. Genes are also presented overlayed in Figure 8.

**Supplemental Figure 9:**
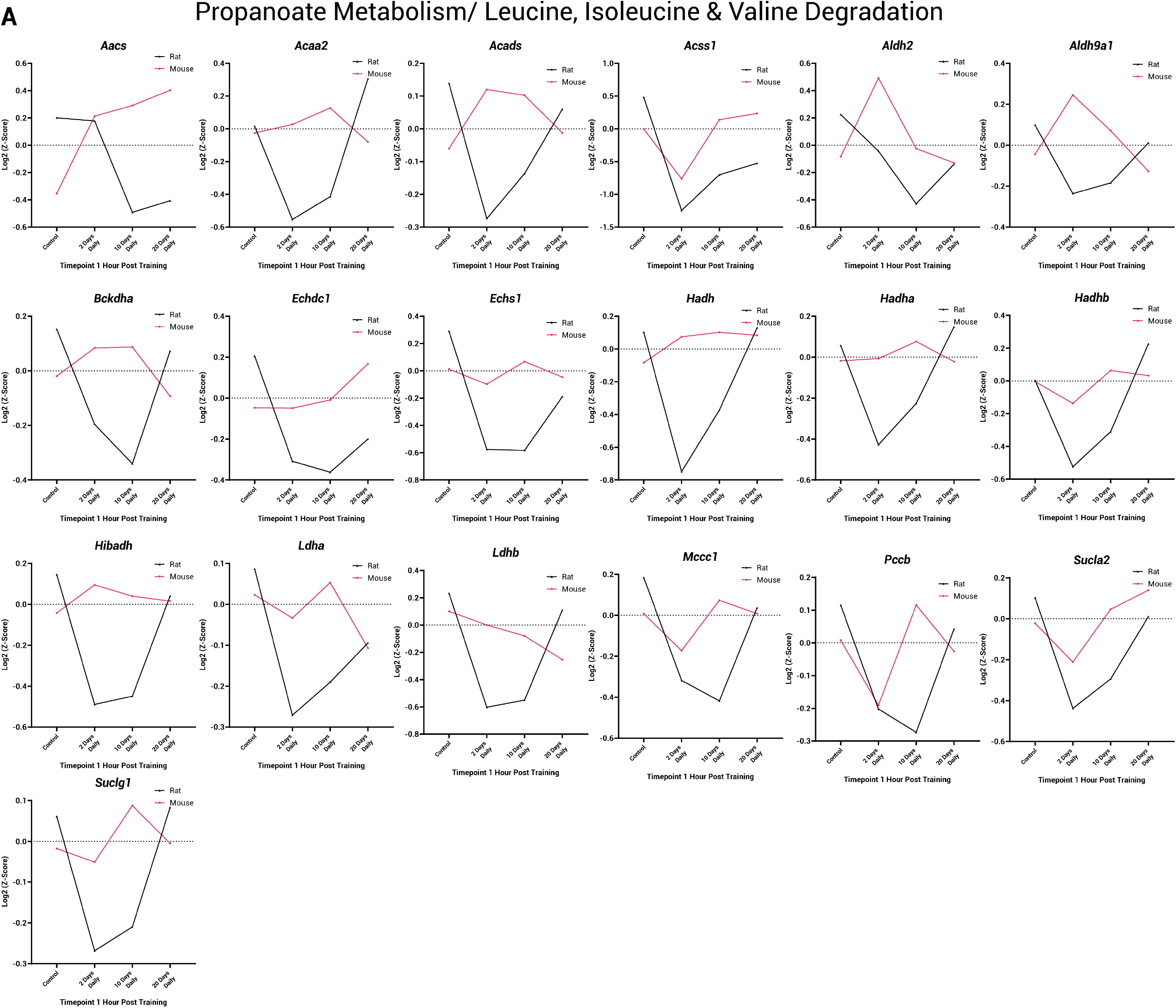
(A) Individual gene plots for genes associated with Propanoate Metabolism and Leucine, Isoleucine and Valine Degradation. Rat gene expression is indicated by black lines, mouse gene expression is indicated by pink lines. Genes are also presented overlayed in Figure 8.

